# Translational toolkit for reproducible, cross-study profiling of human ageing hallmarks in human blood and tissue

**DOI:** 10.64898/2026.04.20.719545

**Authors:** Kristina Tomkova, Jonathan W. Lewis, Thomas Nicholson, Animesh Acharjee, Aidan Anderson, Jonathan P. Barlow, Kathryn Frost, Thomas Jackson, João Pedro de Magalhães, Sudip Mondal, Amy J. Naylor, Aline Nixon, Nicholas Rattray, Paula Rudzinska, Amanda V. Sardeli, Claire J. Steves, Carly Welch, Daisy Wilson, Niharika A. Duggal, Jose R. Hombrebueno, Simon W. Jones, Helen M. McGettrick

**Affiliations:** Department of Inflammation and Ageing, School of Infection, Inflammation and Immunology, College of Medicine and Health, University of Birmingham, Edgbaston, Birmingham, B15 2TT; NIHR Birmingham Biomedical Research Centre, University Hospital Birmingham and University of Birmingham, Birmingham, UK; Department of Cancer and Genomic Sciences, University of Birmingham, Birmingham, UK; Cellular Health and Metabolism Facility School of Sport, Exercise and Rehabilitation Sciences University of Birmingham, Birmingham, UK; Strathclyde Centre for Molecular Bioscience, University of Strathclyde, Glasgow, UK; Department of Twin Research, Kings College London, St Thomas’ Hospital Campus, 3 Westminster Bridge Road, London SE1 7EH

**Keywords:** Biology of ageing, hallmarks of ageing, optimised methodology

## Abstract

**Background:** Ageing is a complex, multi-dimensional process, underpinned by interacting biological hallmarks that collectively contribute to functional decline and increased susceptibility to disease. While considerable progress has been made in delineating individual ageing pathways, translation into human studies has been hindered by methodological heterogeneity and a lack of standardised, multi-system approaches. Here, we describe a validated, high-resolution toolkit for the simultaneous quantification of multiple ageing hallmarks in clinically accessible human samples, encompassing cellular senescence, immune ageing, inflammation, mitochondrial function, mTOR signalling, autophagy, genomic instability, and stem cell exhaustion.

**Methods:** Blood (25ml) was obtained from young and aged donors (26–81y). Deep immunophenotyping was performed using a novel 30-colour spectral flow cytometry panel. T-cell mTOR activation and autophagic flux were assessed by flow cytometry. Metabolic flux was measured by Seahorse. From whole blood (4 ml), muscle, and adipose tissue (AT) (obtained during elective hip arthroplasty) RNA, DNA, AT stem cells, and myoblasts were isolated. DNA copy number and senescent cell burden were assessed by q-PCR and SA-β-gal staining, respectively.

**Findings:** Utilising this toolkit, we identified pronounced age-related immune remodelling, increased senescent T-cell burden, diminished mitochondrial capacity and altered mTOR–autophagy signalling between healthy young and aged donors. Furthermore, metabolism was significantly affected by anti-coagulant and freezing sample before analysis.

**Interpretation:** This integrated platform provides a foundation for reproducible, cross-study analyses and facilitates translational investigation of interventions targeting health-span extension.

**Funding:** Wellcome Leap Dynamic Resilience program (co-funded by Temasek Trust).

## Introduction

Global increases in lifespan have not been matched by equivalent gains in health-span, leading to an increasing prevalence of multimorbidity and age-related disease, with major socioeconomic and healthcare impacts^1^. Slowing biological ageing is therefore a central priority for developing novel interventions that extend health-span.

The identification of the Hallmarks of Ageing – a set of twelve interconnected biological processes including cellular senescence, mitochondrial dysfunction and chronic inflammation that contribute to age-related decline - has accelerated progress in the field^2^. Despite extensive preclinical data, translation is hindered by two major challenges: First, hallmarks are often studied in isolation, obscuring their relative and integrated contributions to clinical phenotypes, such as frailty and dementia^3^. Secondly, methodological heterogeneity across studies (e.g., sample handling, analytical protocols, and experimental conditions) limits reproducibility and comparability^4–6^.

Standardised, validated approaches for measuring ageing hallmarks in human samples are urgently needed to define robust biomarkers of biological age, enable integrative analyses across datasets, and support clinical evaluation of geroprotective interventions. Here, we describe a validated methodological toolkit for simultaneously quantifying multiple hallmarks of ageing in clinically accessible quantities of blood and peripheral tissue (**Fig. 1**). We detail sample requirements, compare commonly used techniques and demonstrate reproducible detection of hallmark differences between young and aged donors. This toolkit establishes a foundation for standardised, cross-study assessment of ageing biology and will facilitate translational research to extend health-span.

**Figure 1.**
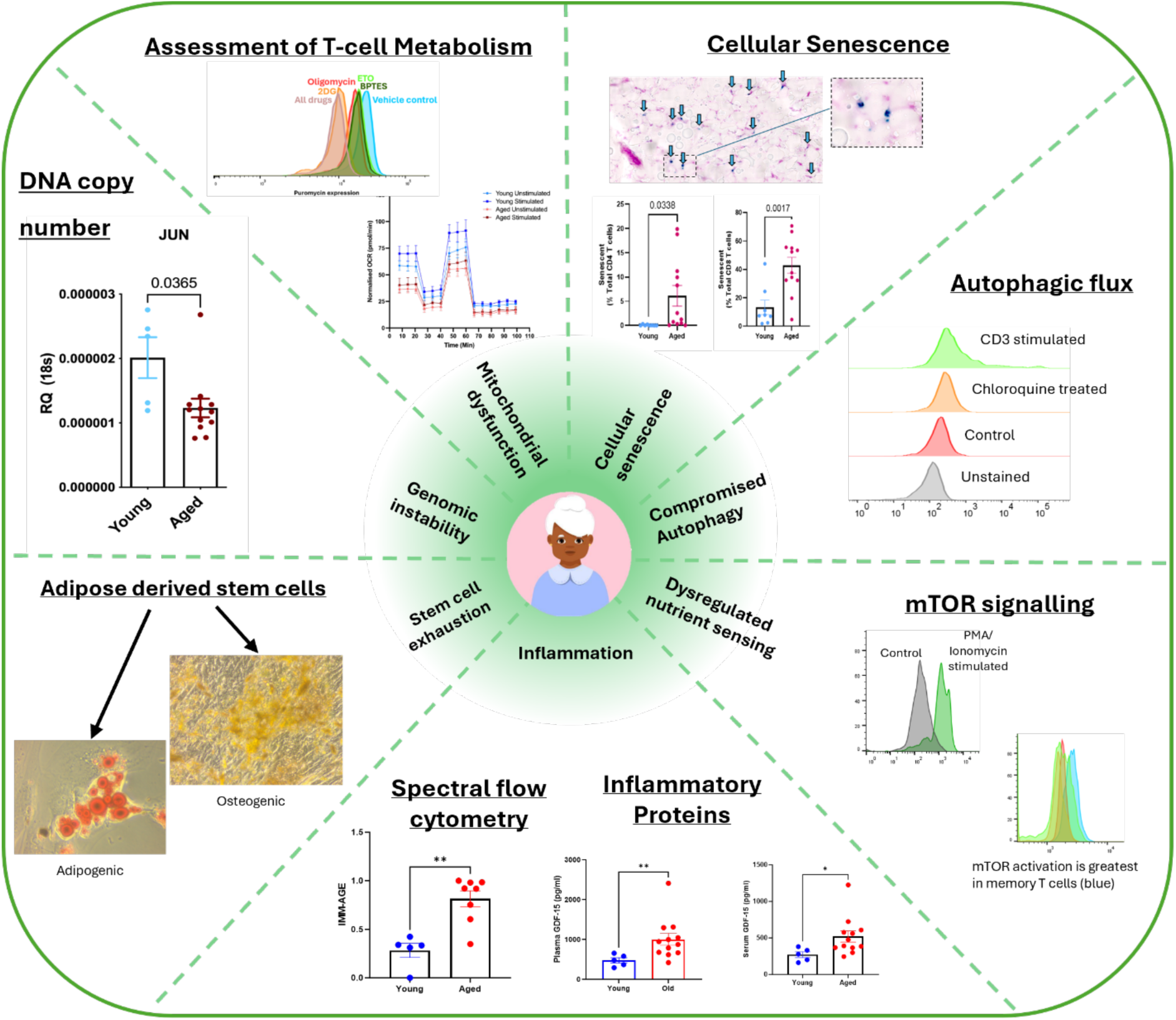
Ageing toolkit summary. Schematic overview of all validated methods to enable the comprehensive profiling of ageing hallmarks (from top right clockwise - cellular senescence, compromised autophagy, dysregulated nutrient sensing, inflammation, stem cell exhaustion, genomic instability, mitochondrial dysfunction) across clinically feasible blood and adipose tissue samples.

## Results

### High-resolution multi-system profiling of cellular senescence

The accumulation of senescent cells in tissues is considered a driver of ageing, and their targeted removal in mice can prevent a broad range of age-related diseases^7^. Senescent cells secrete numerous pro-inflammatory cytokines and other signalling molecules, collectively named the senescence-associated secretory phenotype or SASP, which can further impair function and induce senescence in neighbouring cells in a paracrine fashion^2,8^. Despite extensive research, there is no standardised approach to quantifying and reporting cellular senescence: various measurements can be used, including intracellular markers of cell cycle arrest (e.g., p16, p21, p53), DNA damage (e.g., γ-H2AX), nuclear membrane proteins (e.g. lamin B1), alterations in cellular morphology/biophysical properties, and senescence-associated β-galactosidase (SA-β-gal) activity^9^. Furthermore, studies often assess only a single marker or biological process to confirm senescence, compounding the lack of standardisation and failing to fully capture the burden of cellular senescence within and across tissues/organ systems^9^. To enable effective comparison of senescence phenotype across geroscience studies, wider integration of high throughput and high-resolution techniques is required.

To address this, we validated a novel spectral cytometry panel enabling quantification of circulating leukocyte subpopulations and integration of intracellular senescence markers (p16 and p21), as well as IMM-AGE score^10,11^ calculation using 3×10^5^ peripheral blood mononuclear cells (PBMCs) **(Fig 2A, Supp. Table 1, 2)**. To control for methodological variability, we compared cryopreserved versus freshly isolated PBMCs and evaluated the effect of different anticoagulants (EDTA vs heparin). Neither influenced the distribution of immune populations (**Supp. Fig. 2A, B)**^12^. Notably this represents a unique comparison of the impact of anticoagulant type on immune cell populations. Importantly, our panel detected a robust upregulation of p16 and p21 following irradiation-induced senescence, confirming its ability to identify intracellular senescence markers **(Supp. Fig. 2C, D).**

**Figure 2.**
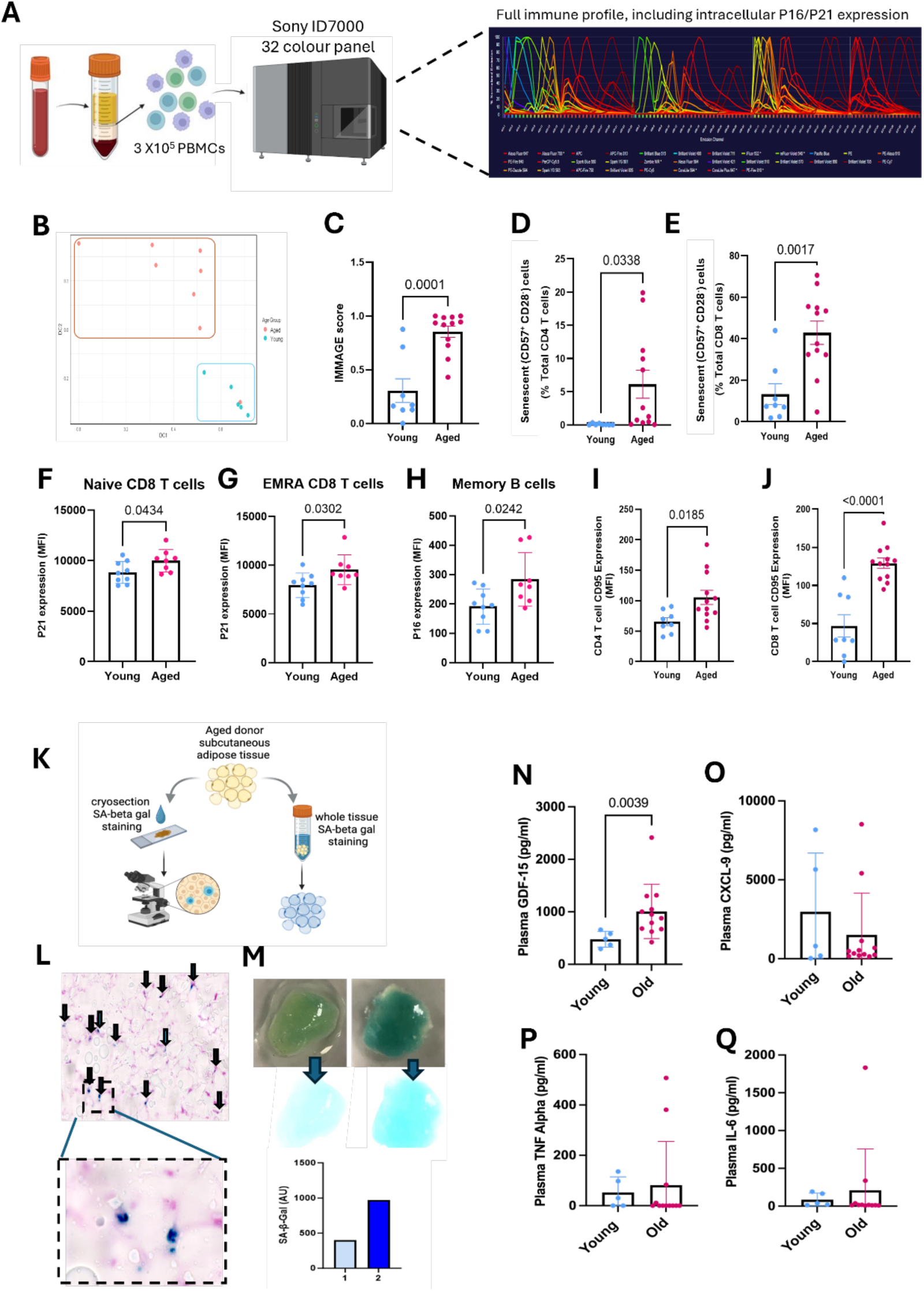
Integrated profiling of immunosenescence, tissue senescence, and circulating SASP in young and aged individuals. **A.** Schematic overview of a novel 32-color spectral flow cytometry panel for profiling circulating immune populations and intracellular senescence markers p16 and p21. **B.** PCA analysis of young and aged peripheral blood mononuclear cells (PBMCs), integrating all identified immune cell subsets. **C.** IMMAge score calculated based on PBMC subsets for young and aged participants in young (n=8) and aged (n=12) participants. **D.** CD4+ CD57+CD28- and **E.** CD8+ CD57+CD28-senescent cells expressed as a percentage of total CD4+ or CD8 T cells in young (n=8) and aged (n=12) participants. Intracellular p21 expression in **F.** Naïve and **G.** EMRA CD8+ T cells from (n=9) young and aged (n=12) participants. **H.** Intracellular p16 expression assessed as median fluorescent intensity in Memory B cells (CD19+ CD38− CD24+) from (n=9) young and aged (n=12) participants. Expression of CD95 in **I.** CD4+ and **J.** CD8+ T cells between in young (n=8) and aged (n=12) participants. **K.** Schematic overview of cryosection and whole-tissue approaches for detecting cellular senescence using β-galactosidase staining (SA-beta gal) in human adipose tissue. Representative images of positive β-galactosidase staining (blue) of senescent cells in **L.** 20 µM human adipose tissue cryosections and **M.** whole human adipose tissue following 24h incubation. Expression of SASP markers **N.** GDF15, **O.** CXCL-9, **P.** TNF alpha, and **Q.** IL-6 in plasma samples from young (n=5) and aged (n=12) participants, determined by ELISA. Data are mean ± S.E.M. Statistical significance was assessed (C-F, H-J) unpaired t-tests or (G, N-Q) Mann Whitney U.

Applying this panel to assess age-associated immune changes, older adults (donor demographics presented in **Supp Table 3**) exhibited a significant reduction in B cells and an increase in dendritic cells compared to young donors **(Supp. Fig. 3A)**, with no significant differences in T cells, monocytes, NK cells, or circulating endothelial cells **(Supp. Fig. 3A)**. Subset-level analyses revealed marked compositional shifts with age, as shown by principal component analysis **(Fig. 2B)**. Consistent with previous reports^13–15^, aged participants had significantly higher frequencies of central memory T-cells, effector memory and EMRA CD8⁺ T cells, and non-classical (CD14⁺CD16⁺) monocytes, and fewer naïve CD4⁺ and CD8⁺ T cells **(Supp. Fig. 3B)**. IMM-AGE scores^10^ confirmed significant immune ageing in the older cohort **(Fig. 2C).**

**Figure 3.**
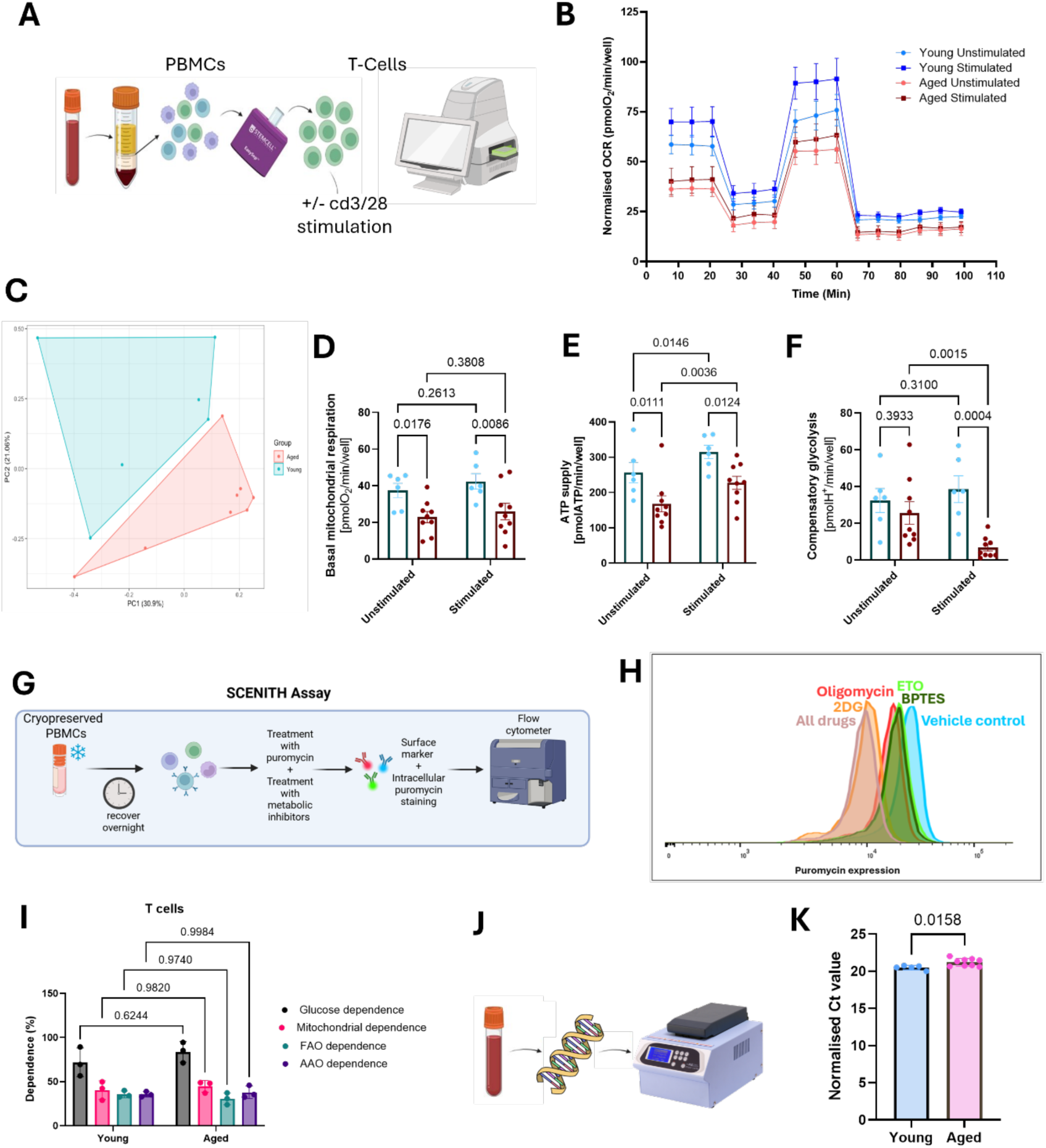
Assessment of immunometabolic function with ageing. **A.** Experimental approach used to measure mitochondrial function in isolated T cells using the Seahorse XF platform. **B.** Representative OCR (oxygen consumption rate) traces from young and aged T cells with or without stimulation with CD3/CD28 for 24 hours. **C.** PCA analysis of overall mitochondrial function in T-Cells isolated from young (n=6) and aged (n=9) aged participants. **D.** Basal Mitochondrial function, **E.** ATP production rate and **F.** compensatory glycolysis in T-cells isolated from (n=6) and aged (n=9) aged participants. **G**. Schematic overview of the SCENITH assay and **H.** representative metabolic shifts observed in peripheral blood mononuclear cells (PBMCs) following treatment with pathway-specific inhibitors (AAO: Amino acid oxidation, BPTES: Bis-2-(5-phenylacetamido-1,3,4-thiadiazol-2-yl)ethyl sulphide, ETO: Etoxomir). **I.** Metabolic dependencies in CD3+ T Cells From young (n=3) and aged (n=3) participants. **J.** Schematic overview of whole blood mitochondrial DNA copy number assessment by RT-qPCR, comparing mitochondrial DNA to nuclear DNA levels. **K.** Mitochondrial DNA copy number in young (n=6) and aged (n=9) participants. Data are mean ± S.E.M. Statistical significance was assessed (**K**) unpaired t-tests and (**D-F, I**) Sidak’s post-test.

Older adults also displayed significantly elevated frequencies of senescent (CD57⁺CD28⁻) CD4⁺ and CD8⁺ T cells **(Fig. 2D, E)**, which co-expressed elevated levels of both p21 and p16 (**Supp. Fig 2E, F**). Moreover, higher p21 expression was detected in naïve and EMRA CD8⁺ T cells **(Fig. 2F, G)** and increased p16 levels seen in memory B cells **(Fig. 2H)** in older donors compared to young counterparts. Older donors also exhibited significantly more apoptotic (CD95⁺) CD4⁺ and CD8⁺ T cells **(Fig. 2I, J)**. Collectively, this panel provides a scalable, clinically relevant platform for high-resolution assessment of immune ageing in human blood, offering the potential to support early detection of accelerated ageing and longitudinal monitoring of therapeutic efficacy.

Beyond immunosenescence, tissue-resident senescent cells contribute to age-related tissue damage and impaired repair^16^. Few studies measure senescence across multiple systems in the same individual^17,18^. To encourage a more holistic approach to ageing research, we optimised two methods of examining senescence in subcutaneous white adipose tissue (SAT) - a depot known to accumulate senescent cells with age^19^ that can be safely accessed through minimally invasive procedures in older adults^20^. We have validated methodologies for the detection of senescent cells using SA-β-gal staining in both frozen adipose tissue sections and fresh whole adipose tissue biopsies **(Fig. 2K)**. Using frozen SAT, positive SA-β-gal staining was observed in 20 μm sections of OCT-embedded sections from aged donors **(Fig. 2L)**, though this approach requires temperature controlled cryosectioning facilities to avoid tissue damage during sectioning^21^. To improve accessibility, we optimised a whole-tissue staining method^22^, where senescent load is quantified after overnight immersion in SA-β-gal solution and imaged analysis using the ImageJ software **(Fig. 2M)**. Strong staining was consistently detected in fresh SAT at pH 5.5 but not pH 7, while fixation prior to staining markedly reduced SA-β-gal signal (**Supp. Fig. 4A**). Together, these methods provide validated options for assessing senescence load in SAT, accommodating variable equipment access and processing constraints, thereby enhancing the toolkit’s flexibility and translational relevance.

**Figure 4.**
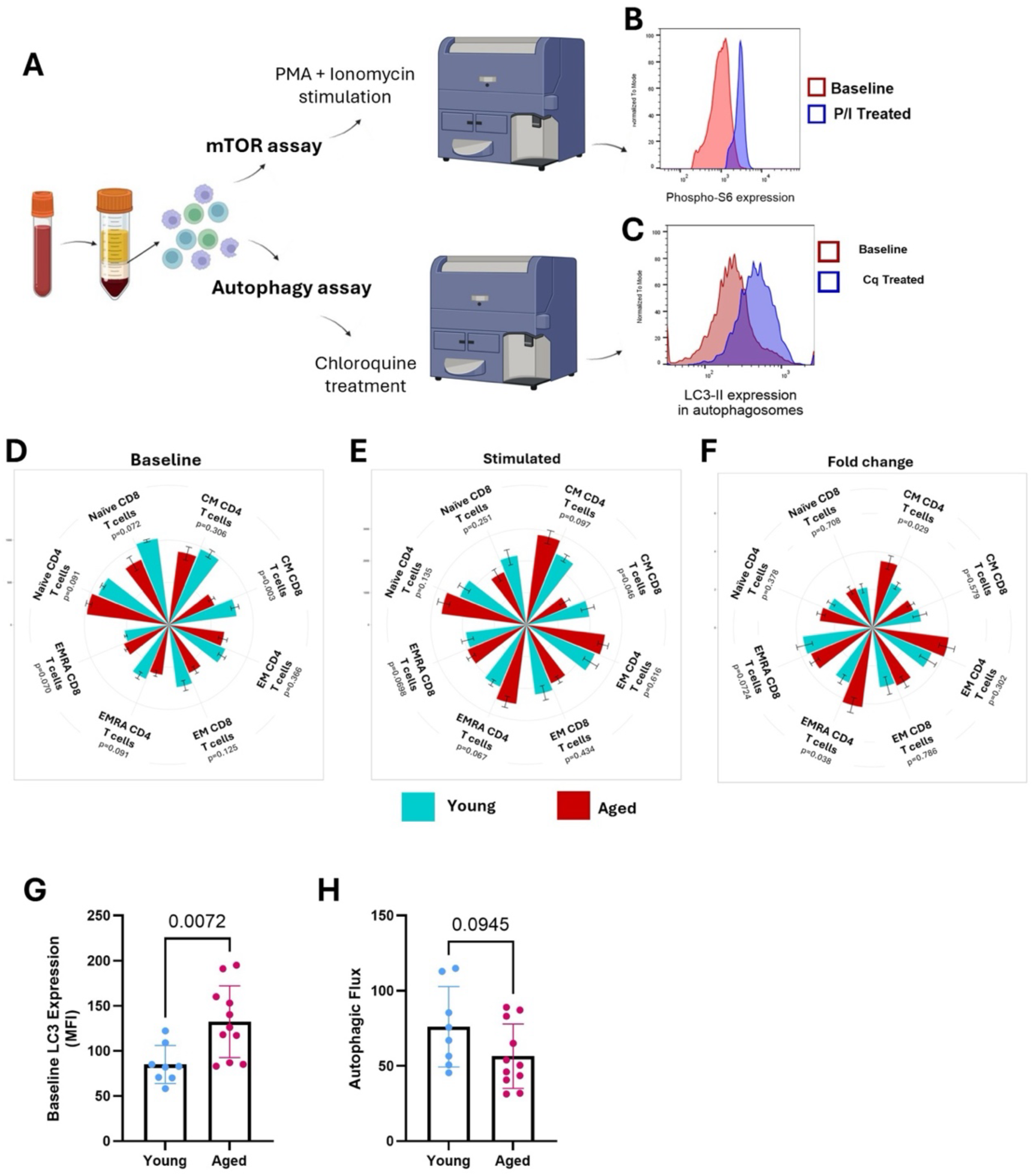
Ageing is associated with dysregulation of functional hallmarks of ageing in immune cells. **A.** Experimental design for assessing nutrient sensing (mTOR activation) and autophagic flux in cryopreserved PBMCs using flow cytometry. **B.** Representative histograms showing phospho-S6 expression in PBMCs at baseline (untreated) and following 30-minute stimulation with PMA (phorbol-12-myristate-13-acetate) and ionomycin. **C.** Representative histogram showing LC3-II (microtubule-associated protein 1A/1B-light chain 3) accumulation in autophagosomes of T cells at baseline and after 2-hour treatment with 100 µM chloroquine (Cq). Radar plots depicting phospho-S6 expression assessed as median fluorescence intensity (MFI) in **(D)** CD4+ and **(E)** CD8+ T cells at baseline and after 30-minute stimulation with PMA and ionomycin in young and aged participants. **F.** Radar plots showing phospho-S6 fold-change relative to young participants in CD4+ and CD8+ T cells following stimulation. **G.** Baseline autophagosomal LC3-II expression in T cells from young (blue, n=6) and aged (red, n=12) participants. **H.** T-cell autophagic flux following 2-hour treatment with 100 µM chloroquine. Data are mean ± S.E.M. Statistical significance was assessed (D-H) unpaired t-tests.

Circulating SASP components in plasma or serum are frequently used as surrogate systemic indicators of senescence^9^. Given its highly complex composition, most studies measure a limited cytokine panel, commonly IL-6 and TNFα^23^. Recently, CXCL-9 (MIG) and GDF-15 have emerged as novel serum SASP components that positively correlate with accelerated biological ageing, age-related disease and mortality^24^. Here, GDF-15 levels were significantly higher in healthy aged versus young participants in both plasma (**Fig. 2N**) and serum (**Supp. Fig. 4B**), whereas CXCL-9, TNFα and IL-6 showed no age-related differences (**Fig. 2O-Q, Supp. Fig. 4C-E**). To highlight the important differences in SASP quantification and interpretation between serum and plasma, we compared absolute concentration of the SASP molecules in both sample types. GDF-15 levels were significantly higher in plasma, whilst CXCL9 and IL-6 levels were significantly lower compared to matched serum (**Supp. Fig. 4F-I**). These findings demonstrate that plasma and serum are not interchangeable for SASP analysis and underscore the importance of consistent sample selection for accurate interpretation.

Overall, this toolkit unifies optimised and validated methods for assessing senescence across diverse sample types, providing the first integrated framework for multi-system analysis of human ageing biology. By accommodating varying equipment availability and processing constraints, it offers practical accessibility for both research and clinical laboratories. Applying these complementary assays reveals that senescence is not uniformly manifested across tissues or cell types, highlighting the importance of multimodal, cross-tissue profiling to capture the heterogeneity of senescence load *in vivo*.

### Multi-dimensional profiling reveals age-associated mitochondrial decline in human immune cells

Mitochondrial dysfunction is a central hallmark of ageing, characterised by impaired energy production, reduced metabolic flexibility, altered substrate utilisation^25^, and the accumulation of mitochondrial DNA (mtDNA) mutations and copy number alterations^26^. PBMCs are widely used to study mitochondrial function, but no consensus exists on optimal methodologies, which vary in assay design and readouts. For example, metabolic flux analyses may differ in substrate availability, cell handling and assay endpoints^27–29^, while mtDNA quantification is inconsistently reported as copy number, deletions or mutational burdens^30^. Without methodological clarity, it remains difficult to discern whether metabolic differences across cohorts reflect a true ageing effect or are due to technical artefact, limiting the identification of robust metabolic biomarkers of human ageing. To overcome this, we systematically optimised assays to assess mitochondrial metabolic function and quantify mtDNA copy number.

The Agilent Seahorse platform is the current gold-standard method to evaluate metabolic flux in vitro; however, differences in assay media composition, cell seeding density, drug concentrations and incubation parameters complicate cross-study comparisons^27–29^. We employed a modified Seahorse XF Mito Stress Test to measure oxygen consumption rate (OCR) and extracellular acidification rate (ECAR) in T cells under basal conditions and following CD3/CD28 stimulation, which mimics T cell receptor engagement (**Fig. 3A**). T cells were chosen because their function relies heavily on metabolic plasticity, and age-associated metabolic rewiring in these cells has been linked to impaired vaccine responses, chronic inflammation, and tissue repair^31^. To ensure broad reproducibility, we used generic mitochondrial stress test compounds (oligomycin, BAM15, rotenone/antimycin A, and monensin) rather than pre-formulated Seahorse XF kits, allowing precise control over drug concentrations and utilisation across laboratories.

Seeding density was optimised using freshly isolated T cells rested 16 h overnight to ensure consistent results within the dynamic range of the Seahorse XF. Densities <200k cells/well yielded poor responses, while 200–300k T-cells/well produced robust, reproducible OCR and ECAR responses (**Supp. Fig. 5A–C**). We recommend 200k cells/well to balance sufficient signal-to-noise and greater delta in maximal respiration/spare respiratory capacity, whist avoiding oxygen or nutrient limitations associated with overcrowding of wells with >300k. Critically, cryopreserved T cells recovered for 16 h prior to assay could not replicate these responses these results (**Supp. Fig. 5D-E**), indicating consistent metabolic flux requires the use of fresh samples, which may not be feasible in some settings.

**Figure 5.**
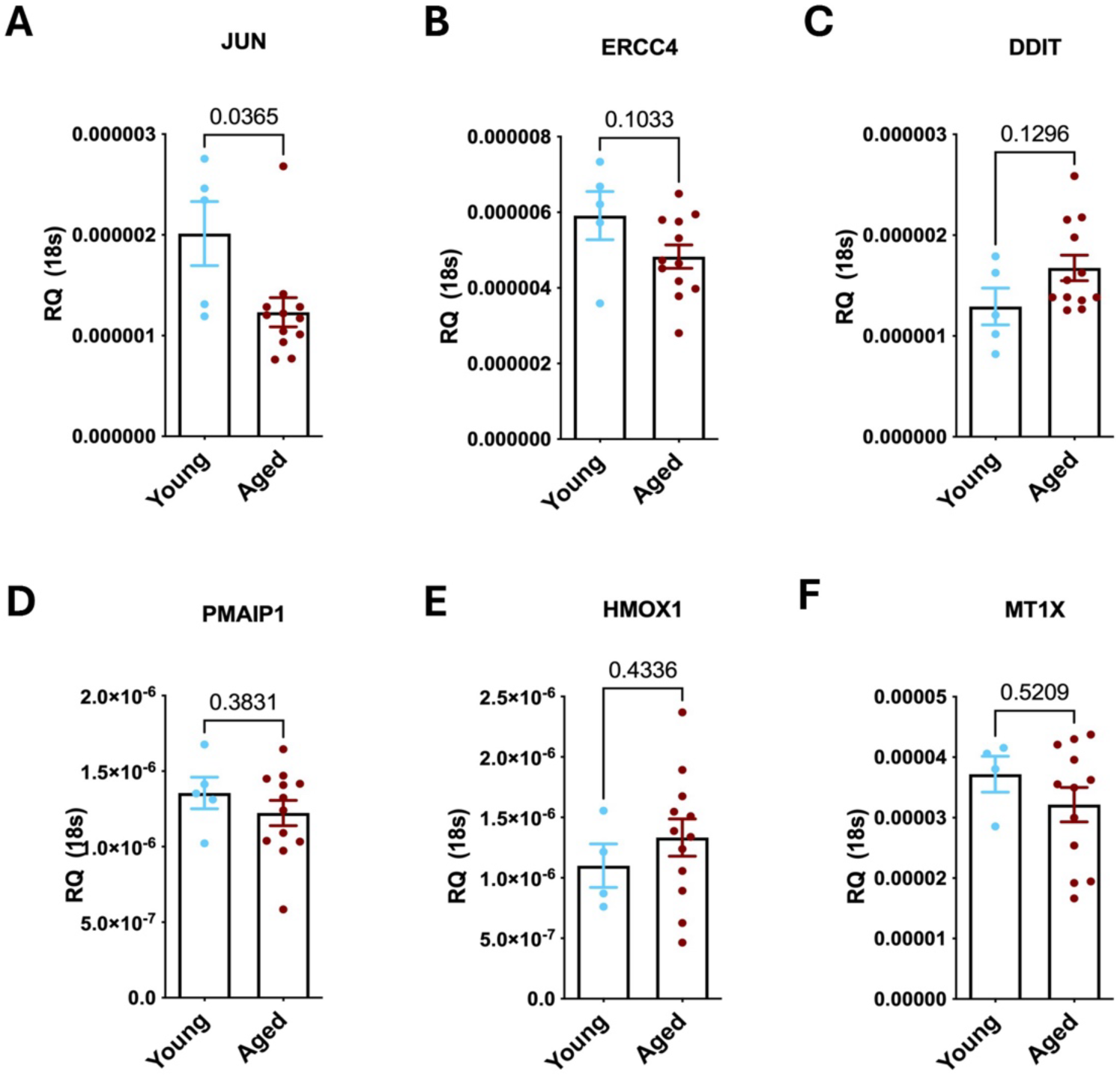
Age-Driven Genomic Instability in Human Blood. mRNA expression of **(A)** *JUN,* **(B)** *ERCC4,* **(C)** *DDIT,* **(D)** *PMAIP1,* **(E)** *HMOX1* and **(F)** *MT1X* in whole blood from young (blue, n=5) and aged (red, n=12) participants determined by q-rtPCR. Data are expressed as the relative quantification (RQ) compared to *18s.* Data are mean ± S.E.M. Statistical significance was assessed (A-B, D-F) unpaired t-tests or (C) Mann Whitney U.

Next, we examined the impact of anticoagulant (heparin, EDTA) on T cell metabolic flux – an often overlooked variable known to influence immune cell mitochondrial function^32^. Basal OCR was similar between anticoagulants, but EDTA-derived T cells exhibited higher OCR after BAM15 injection compared with T cells from heparinised blood (**Supp. Fig. 5F**), reflecting increased maximal respiratory capacity. Furthermore, EDTA-derived T cells displayed greater basal OCR following CD3/CD28 stimulation, which was further amplified following BAM15 injection (**Supp. Fig. 5F**), suggesting EDTA better preserves the dynamic respiratory range of T cells under activation conditions. In contrast, ECAR responses at baseline and after activation were unaffected by anticoagulant choice (**Supp. Fig. 5G**), suggesting glycolytic capacity is not influenced by type of anticoagulant. Collectively, these data indicate that metabolic flux analyses in T cells are most robust using freshly isolated PBMCs from EDTA-treated blood.

Using this optimised protocol, T cells from aged donors exhibited distinct metabolic alterations compared to young donors. PCA analysis and Seahorse OCR traces revealed impaired mitochondrial output (**Fig. 3B, C**), with the greatest deficits following BAM15-induced uncoupling, indicating reduced mitochondrial reserve in aged T cells (**Fig. 3B**). Aged T cells also showed significantly reduced basal OCR and ATP supply rates at rest and after CD3/CD28 stimulation (**Fig. 3D-E**), along with blunted compensatory glycolysis in response to monensin after stimulation (**Fig. 3F**), reflecting reduced metabolic flexibility. Comprehensive profiling of oxidative and glycolytic contributions to ATP supply further confirmed this age-associated decline in metabolic adaptability, including significantly reduced phosphorylation linked respiration, maximal respiration, and non-mitochondrial respiration after stimulation, as well as reduced mitochondrial ATP supply in both baseline and stimulated states (**Supp. Fig. 6**). Together, these results demonstrate a reproducible, physiologically relevant protocol for assessing age-associated immunometabolic decline.

**Figure 6.**
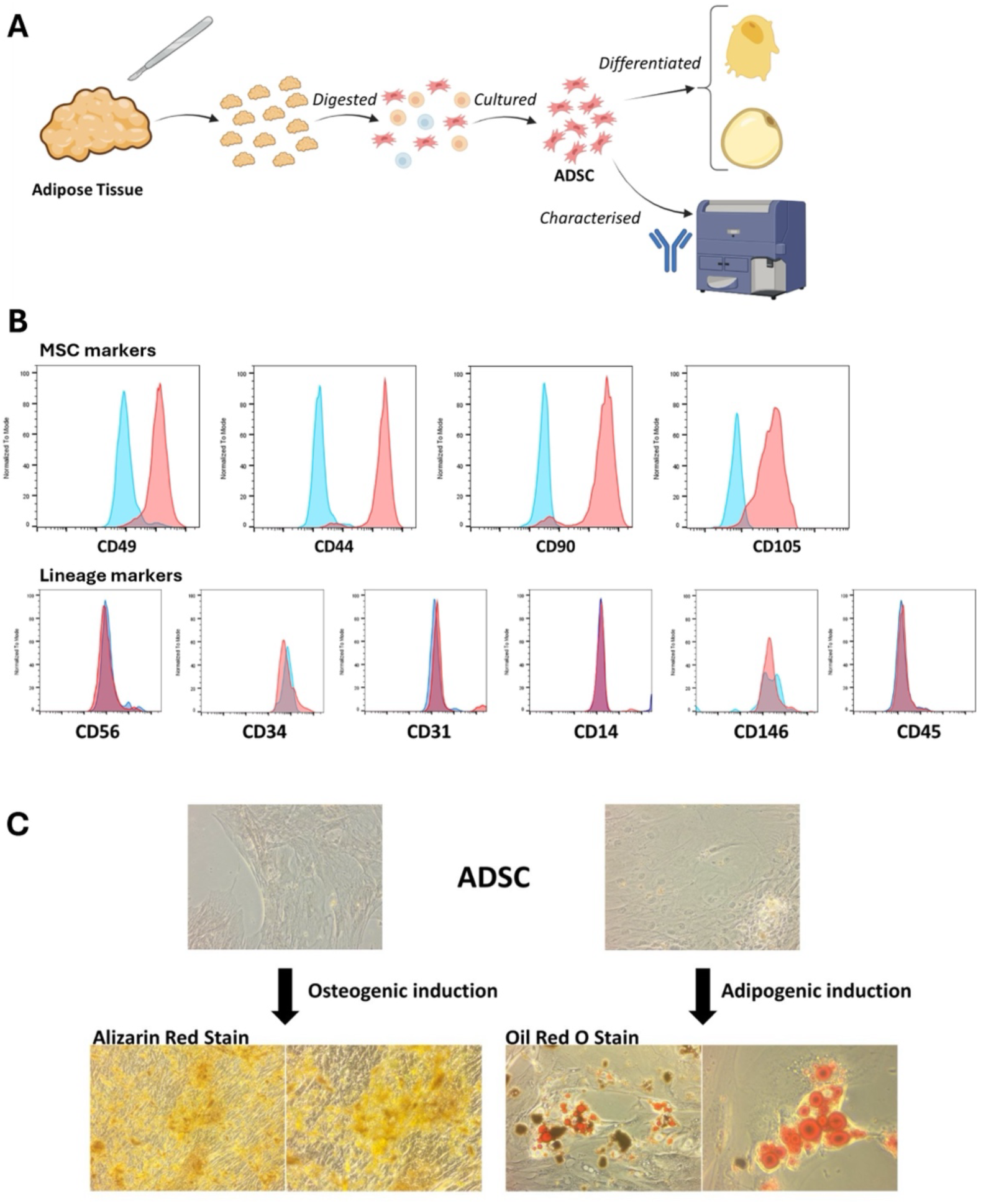
Human adipose-derived precursor cells provide a model to study stem cell exhaustion with age. **A.** Schematic representation of the experimental design demonstrating, collagenase mediated isolation of adipose derived stem cells (ADSCs) from human subcutaneous adipose tissue, derived from aged participants. Stemness of isolated cells was determined by assessment of differentiation capacity and positive and negative expression of stem cell and lineage markers respectively. **B.** Representative histograms demonstrating isolated ADSCs expressed mesenchymal stem cells: CD49, CD44, CD90 and CD105 and were negative for lineage markers: CD56, CD34, CD31, CD14, CD146 and CD45. **C.** Left: Representative images demonstrating positive staining of (Oil Red O stained lipid droplets) following adipogenic differentiation. Right: Representative images of osteogenic positive Alizarin red stained mineral, following osteogenic differentiation.

While Seahorse flux analysis provides dynamic information on population-level responses, its reliance on freshly isolated cells and specialised equipment limits its application in large-scale or multicentre ageing studies. To address this, we validated SCENITH^33^, a flow cytometry-based method for single-cell metabolic profiling that quantifies cellular dependencies on glycolysis, oxidative phosphorylation, fatty acid oxidation, and amino acid oxidation. SCENITH couples metabolic challenge induced by specific inhibitors with immune cell subset analysis, allowing parallel assessment of metabolism and immune phenotype at single-cell resolution (**Fig. 3G, H**). Utilising PBMCs from young donors, SCENITH reliably demonstrates the expected reductions in puromycin incorporation following oligomycin or 2-DG treatment in both fresh and cryopreserved cells (**Supp. Fig. 7A**). As seen with Seahorse, EDTA-treated blood generated more consistent metabolic shifts than heparinised blood (**Supp. Fig. 7B**).

Applying SCENITH to PBMCs from young and aged donors collected in EDTA, we have not observed any significant age-related differences in total CD3⁺ T cells or across the T cell subsets (**Fig. 3I, Supp. Fig. 7C–J**), likely due to insufficient sample size. However, compared to the Seahorse assay, the strength of SCENITH lies in its capacity for high-dimensional metabolic phenotyping through concurrent immunostaining. Combining CD4, CD8, CCR7, and CD45RA markers allowed assessment of metabolic profiles for naïve, central memory (CM), effector memory (EM) and terminally differentiated effector memory (EMRA) T cell subsets in frozen cells (**Supp. Fig. 7C–J**).

Mitochondrial DNA copy number provides a complementary, structural measure of mitochondrial health, distinct from functional assessment by Seahorse or SCENITH. Previous studies often a single mtDNA locus (e.g., ND1, COX1, or the D-loop/control region), with inconsistencies in normalisation approaches (e.g., to a single nuclear gene, total DNA, or cell number) limiting sensitivity and reproducibility. We optimised a qPCR assay using whole-blood genomic DNA^34^ (**Fig. 3J**), with four primer sets targeting distinct mitochondrial regions: mtControl (D-loop), mtND4, mtRNR2, and mtCO1, and normalisation against four nuclear genes (B2M, ACTB, GAPDH, TFRC) to improve accuracy by reducing bias from regional deletions or sequence variability and correcting for cell number and genomic DNA content (**Supp. Fig. 7K, L**). This approached revealed a significant reduction in relative mtDNA copy number in aged compared with young donors (**Fig. 3K**) as demonstrated by increase in Ct values, consistent with age-associated mitochondrial decline observed in published literature^35^.

Together with our optimised Seahorse and SCENITH workflows, these approaches enable multi-dimensional assessment of mitochondrial health, integrating functional flux, single-cell metabolic dependency, and mitochondrial genomic integrity in the same donors. Combining these readouts with senescence and immune phenotyping in matched samples allows direct evaluation of mitochondrial health alongside age-associated immune dysfunction.

### Ageing is associated with dysregulation of mTOR signalling and autophagic flux in human immune cells

The mechanistic target of rapamycin (mTOR) pathway regulates metabolism, protein synthesis, and cellular ageing. Hyperactivation of mTOR signalling has been implicated in driving age-related functional decline, while prolonged inhibition with rapamycin extends lifespan and health-span in model organisms^36^. Despite this, evidence of age-associated alterations in mTOR activity within human immune cells remains limited and is inconsistently reported. Whilst some studies showed that mTOR inhibition within the immune system improves viral immunity^37^ and attenuates autoimmune diseases^38,39^, other studies failed to see any positive effects on the immune system health^40^. Some of these discrepancies can be attributed to the heterogeneity in immune subsets measured and methodological difference^41^.

To address this, we optimised a flow cytometry-based assay to quantify the levels of ribosomal protein S6 kinase (S6K) phosphorylation, a canonical downstream target of mTOR, in PBMCs from young and aged donors (**Fig. 4A**). To examine different immune cell states, we assessed unstimulated and Phorbol 12-Myristate 13-Acetate (PMA)/Ionomycin stimulated PBMCs^11,42,43^, which elicited a robust induction of S6K phosphorylation (**Fig. 4B**). Assay performance was unaffected by sample cryopreservation or anticoagulant type (**Supp. Fig. 8A-D**). The inclusion of surface marker staining enabled analysis of mTOR activity across different immune subsets. Unstimulated central memory (CM) CD8^+^ T cells from aged donors showed significantly lower phosphorylation of S6K compared to young, a difference maintained after stimulation (**Fig. 4D-E**). In contrast, CM and EMRA CD4⁺ T cells showed an age-related hypersensitivity to stimulation, exhibiting higher mTOR activity compared to young donors (**Fig. 4F**). Whilst an age-related decline in S6K phosphorylation was also seen in other unstimulated T-cell subsets (**Fig. 4D-F)**, this did not reach statistical significance. These results highlight subset-specific differences in mTOR activity and the value of high-resolution approaches.

Regulated in part by mTOR signalling, autophagy is a fundamental process required for maintenance of protein and organelle quality. Ageing is associated with dysregulated autophagy, in multiple cell types, leading to impaired clearance of intracellular macromolecules and the accumulation of dysfunctional organelles^44^. Despite this central role in cellular homeostasis and ageing, measuring autophagic flux in primary human immune cells is challenging due to its highly dynamic nature and often requires live-cell imaging or pharmacological manipulation^45^. Consequently, many studies rely on indirect or surrogate markers, which can fail to capture autophagic flux or cell-type differences^45^.

We established a flow cytometry assay in T cells to quantify autophagic flux using the lysosomal degradation inhibitor chloroquine and autophagosome-specific LC3II antibodies^43^ (**Fig. 4A, C**). Dose-response testing revealed maximal LC3II accumulation at 100 µM chloroquine after 2h (**Supp. Fig. 8E**). Cryopreserved T cells displayed higher dead-cell proportions when analysed immediately, but overnight resting restored LC3II expression and viability to levels comparable with fresh cells (**Supp. Fig. 8F–I**). Anticoagulant choice had no significant effect (**Supp. Fig. 8J–K**). To improve accessibility, we optimised independent reagents: 0.1% saponin effectively permeabilised membranes comparable to Reagent B provided in the CYTEK Guava® Autophagy kit, removing non-autophagosome LC3II (**Supp. Fig. 8L**). Of note, lower concentrations of saponin resulted in incomplete permeabilization, whilst higher concentrates caused cell death (**Supp. Fig. 8L**). Similarly, an alternative commercially available LC3II antibody performed comparably to the CYTEK Guava® kit antibody (**Supp. Fig. 8M**). Using this assay, we observed an age-related significant increase in LC3II expression in untreated T cells (**Fig. 4G**), with a trend towards decreased autophagic flux (**Fig. 4H**) compared to young donors, suggesting an impaired autophagosome clearance. This kit-independent, flow cytometry-based approach delivers a sensitive and reproducible measure of autophagic flux in human T cells, overcoming accessibility barriers.

Together, these optimised assays provide robust, flow cytometry-based measurement to reliably quantify of mTOR activity and autophagic flux in cryopreserved human immune cells. Integrated with our senescence and metabolic profiling, this platform would enable a comprehensive assessment of functional ageing hallmarks and could offer a foundation for evaluating geroprotective interventions targeting nutrient sensing and proteostasis in humans.

### Minimising pre-analytical variability to assess genomic instability in ageing

Genomic instability, encompassing telomere attrition, accumulated DNA damage, impaired DNA repair, and chromosomal abnormalities, is a primary hallmark of ageing that is thought to drive both tissue dysfunction and the onset of age-related disease^2^. Unlike functional hallmarks, genomic instability is directly measurable at the level of DNA and RNA, yet reliable assessment in human samples has been hindered by variability in extraction methods, storage conditions and marker selection. Factors such as blood collection tube type, storage temperature and duration, RNA/DNA extraction method and reference gene selection can all influence nucleic acid yield, integrity, and downstream quantification^46,47^. Without standardisation, comparisons across cohorts, timepoints, or laboratories risk reflecting pre-analytical artefacts rather than biological ageing effects.

To enable consistent and reproducible downstream applications (e.g. RNA-sequencing, qPCR, telomere length measurement), we optimised nucleic acid isolation from whole blood, comparing extraction methods, anticoagulant and storage conditions on RNA integrity and yield. RNA yields and quality differed significantly between isolation methods (**Supp Table 4**). From 3 ml blood, Tempus ™ tubes containing stabilising reagents for an immediate RNA preservation produced higher RNA yields (63.5 ng/µl; RIN > 7) compared to EDTA blood processed with Qiagen extraction kits (20.3 ng/µl; RIN < 7). Only Tempus-extracted RNA met the RIN ≥7 threshold for RNA-seq library preparation (e.g., Illumina TruSeq, Oxford Nanopore), ensuring unbiased transcriptome profiling. Cold storage of Tempus tubes for 7 days (4 °C or −80 °C) did not reduce yield or integrity compared to freshly processed samples (**Supp Table 4**), supporting multicentre collection and shipment. In contrast, room temperature storage of Tempus tube™ blood for 7 days led to a marked decrease in RNA integrity (mean RIN, 5.9), emphasising the importance of appropriate collection and stabilisation methods ensure detection of low-abundance transcripts relevant to genomic instability and immune ageing are not overlooked.

To capture age-associated genomic instability in circulating immune cells, we measured mRNA expression of six candidate genes (*JUN, ERCC4, DDIT3, PMAIP1, HMOX1, MT1X)* forming a genomic instability-associated stress response signature^48^. This signature was originally defined through integrative analyses linking DNA damage, oxidative stress, and apoptotic pathways and serves as a reproducible marker of cellular responses to genomic instability in our young and aged donors^48^. JUN expression was 50% lower in aged versus young donors (**Fig. 5A**). Whilst we observed a tendency for lower expression of ERCC4, *PMAIP-1* and *MT1X*, and elevated expression of *DDIT* and *HMOX1* transcripts in older adults compared to young, these were not statistically significant when analysed individually (**Fig. 5B-F**).

These optimised RNA isolation and handling protocols provide a robust framework for integrating genomic instability measures into our multi-hallmark toolkit. The limited transcriptional differences in signature genes indicate that genomic instability markers alone capture only part of age-associated stress in circulating immune cells, highlighting the importance of a multi-dimensional toolkit combining transcriptional, metabolic and mitochondrial readouts to comprehensively assess biological ageing in blood.

### Human adipose-derived precursor cells provide a model to study stem cell exhaustion with age

Stem cell exhaustion, defined as the age-associated decline in stemness and regenerative potential, is a central hallmark of ageing that impairs tissue maintenance, repair and overall resilience^49^. While our earlier analyses focus on PBMCs, direct assessment of tissue-resident progenitors remains challenging due to limited accessibility and low cell yield. Prior work in human tissues shows age-related declines in stemness^50,51^, supporting the inclusion of tissue-resident stem cells in our multi-hallmark analyses.

We optimised isolation and characterisation of adipogenic precursor cells from human SAT samples, providing a tissue-based component to our ageing toolkit. Multipotent precursor cells were isolated from 200 mg SAT using collagenase digestion (**Fig. 6A**). Such adipose tissue biopsies can be obtained via minimally invasive and well tolerated incision or needle sampling, making this approach feasible for use in clinical or research settings and more accessible compared to similar studies^52^. These cells robustly expressed canonical mesenchymal stem cell (MSC) markers (CD49, CD44, CD90, CD105) and lacked lineage-specific marker expression (CD56, CD34, CD31, CD14, CD146, CD45), confirming their precursor identity (**Fig. 6B**). Importantly, they retained functional multipotency, differentiating into both adipogenic and osteogenic lineages as evidenced by oil-red-O and alizarin red staining, respectively (**Fig. 6C**). Incorporating adipose tissue resident precursor cells into our ageing toolkit allows future investigation of how stemness and differentiation capacity change with chronological and biological ageing, providing a more holistic view of systemic and tissue-specific cellular dysfunction.

## Discussion

Although human lifespan has increased worldwide, health span has lagged behind, creating an urgent imperative to understand and intervene in the biological processes of ageing. Despite extensive preclinical data, translation into humans has been limited by methodological heterogeneity, the use of isolated markers, and low-resolution approaches that fail to capture the interactions between hallmarks of ageing. To address these challenges, we developed a standardised, clinically feasible toolkit that enables simultaneous profiling of multiple hallmarks across blood and tissues. By integrating high-dimensional spectral flow cytometry, single-cell metabolic readouts, senescence markers, and mTOR/autophagy assays, our platform captures hallmark-specific changes and their interplay within the same individuals, providing a foundation for system-level studies of human ageing^53–55^.

The validity of this platform is supported by alignment with established ageing signatures. Consistent with prior reports, we observed age-associated expansion of senescent and memory T cell subsets, loss of naïve populations^56^, metabolic inflexibility in aged T cells^25^, and systemic elevations of GDF-15 and IL-6 indicative of inflammageing^57^. Crucially, our systematic evaluation of pre-analytical variables revealed that collection tube type, processing delays, cryopreservation, and RNA extraction method significantly impact RNA integrity, soluble marker measurements, and functional readouts. Such technical artefacts, if unaccounted for, can obscure ageing signatures and hinder cross-study reproducibility - a key consideration as geroprotective interventions, including rapamycin analogues, senolytics, and mitophagy activators, progress toward clinical evaluation^58–60^. By embedding standardised protocols and validated reagents, our toolkit mitigates these challenges, enabling reproducible multi-hallmark datasets across laboratories and cohorts. Minimal sample requirements also make the toolkit suitable for frail, multimorbid populations. These individuals are often excluded from research despite representing the primary targets for geroscience interventions^61^.

A central innovation of our framework is its ability to interrogate the interactions between ageing hallmarks within individuals. Mitochondrial dysfunction promotes DNA damage, triggering p53-dependent senescence pathways and amplifying SASP burden^62^. Similarly, mTOR hyperactivation suppresses autophagy, accelerating accumulation of damaged mitochondria and senescent cells^63,64^. By integrating immune phenotyping, metabolic flux, mitochondrial genomics, senescence markers, and nutrient-sensing/proteostatic assays, our toolkit provides the first validated platform to map these mechanistic interconnections in humans. Beyond descriptive profiling, this approach enables generation of composite ageing indices that reflect biological age trajectories more accurately than single biomarkers alone. These multidimensional scores could stratify individuals by mechanistic ageing subtype, supporting targeted, pathway-specific geroprotective interventions.

Our results also underscore the heterogeneity and cell-type specificity of ageing. Population-level analyses often mask subtle but functionally relevant alterations revealed by high-resolution methods. For instance, bulk T cell metabolic profiling showed only modest age-related shifts, whereas subset-specific analyses revealed marked declines in mitochondrial respiration, ATP supply, and metabolic flexibility within memory T cells. Similarly, mTOR signalling exhibited divergent responses across T cell subsets, with age-related hypo- or hyperactivation depending on the population and stimulation state. These observations highlight the importance of high-dimensional, single-cell approaches for accurately capturing ageing biology.

The inclusion of tissue-resident adipose-derived precursor cells provides an additional, novel dimension. We established a reproducible workflow to isolate multipotent stromal precursors from small subcutaneous biopsies, which retain both adipogenic and osteogenic potential^52^. Incorporating these cells allows parallel assessment of systemic and tissue-specific ageing, addressing a key limitation of previous studies that focus exclusively on circulating immune cells. Given the critical role of stem cell exhaustion in tissue maintenance, repair, and organismal resilience^49,50^, this cross-system approach provides a more holistic view of human ageing, linking cellular dysfunction with potential functional outcomes.

Our mTOR and autophagy assays further expand the toolkit’s functional reach. By quantifying S6K phosphorylation and LC3II accumulation in defined immune subsets, we provide high-resolution, flow-cytometry readouts of two central nutrient-sensing and proteostatic pathways. These assays revealed subset-specific dysregulation in aged donors, including reduced mTOR activity in CD8⁺ central memory T cells and diminished autophagic flux in older T cells, highlighting the interconnected remodelling of metabolism, nutrient sensing, and proteostasis with age. Combined with senescence and metabolic modules, these assays enable mechanistic dissection of how interventions targeting nutrient sensing, mitochondrial function, or proteostasis influence immune ageing.

Beyond its biological insight, the platform’s modular design ensures future adaptability. Each assay can be implemented independently or expanded to include emerging hallmarks and omic layers, such as proteomic, metabolomic, or epigenetic clocks^65,66^. Integration with single-cell transcriptomics and standardised functional metrics of muscle performance, cognition, and resilience will be a crucial next step. Linking molecular hallmarks to physiological function will enable the identification of unified, system-level biomarkers that reflect both biological ageing and health-span.

## Conclusion

In summary, we describe a validated, high-resolution, multi-modal platform for profiling human ageing across blood and tissue compartments. By capturing multiple hallmarks, controlling pre-analytical variability, and integrating functional and molecular readouts, the toolkit enables multi-modal profiling of ageing biology in human samples that will have utility in mechanistic investigations, biomarker discovery, and clinical evaluation of geroprotective interventions. Importantly, it offers a framework for precision geroscience, allowing stratification of individuals by mechanistic subtype, longitudinal monitoring of biological ageing, and rigorous assessment of interventions aimed at extending health-span. Collectively, this approach moves human ageing research beyond single-hallmark studies toward an integrated, systems-level understanding of the biology of ageing.

## Authors contributions

**KT, JWL, and TN**: Data curation, Formal Analysis, Investigation, Methodology, Writing-original draft, Writing-review and editing, Visualisation. **AAnderson, KF, PR, and AN:** Methodology, Writing - Review & Editing. **SM, JPB, and AVS:** Formal Analysis, Methodology, Writing-review and editing**. AN:** Writing - Review & Editing. **CJS** Conceptualization, Funding Acquisition, Writing - Review & Editing. **NAD, SWJ, NR, JRH, AJN, CW, AAcharjee, JPdeM, and DW:** Conceptualization, Resources, Supervision, Project administration, Funding Acquisition, Writing - Review & Editing. **TJ and HMM:** Conceptualization, Resources, Supervision, Project administration, Funding Acquisition, Writing-Original Draft, Writing - Review & Editing.

## Data availability statement

Data are available upon reasonable request. All data relevant to the study are included in the article or uploaded as supplementary information.

## Acknowledgements

The authors acknowledge all participants from the Birmingham 1000 Elder’s Cohort for their participation and donation of blood samples. We also acknowledge study participants, research staff, and orthopaedic surgeons at the Royal Orthopaedic Hospital NHS Foundation Trust (Birmingham) and Russell’s Hall Hospital, Dudley in the donation and collection of subcutaneous adipose tissue samples. Analysis was conducted using equipment funded by the College of Medicine and Health Imaging Suite (RRID:SCR_027108), Flow Cytometry Facility (RRID:SCR_027107) and Genomics Birmingham funded by the University of Birmingham. We also acknowledge the research was carried out at the National Institute for Health and Care Research (NIHR) Birmingham Biomedical Research Centre (BRC). Finally, we acknowledge the help and support provided by Rebecca Birch Operational Manager for the project.

## Ethics Approval Statement

The study was conducted in compliance with the Declaration of Helsinki. All human samples were obtained with written, informed consent and approval from the National Research Ethics Committee (NRES 13/NE/0222), Human Biomaterial Resource Centre (Birmingham, UK), West Midlands and Black Country Research Ethics Committee (07/H1203/57), the South-East Scotland Research Ethics Committee (16/SS/0172), Newcastle and North Tyneside 2 Research Ethics Committee (12/NE/0251) or University of Birmingham Local Ethical Review Committee (ERN12_0079).

## Methods

### Ethics

Informed consent was obtained from all participants. Aged donors were enrolled through the Birmingham 1000 Elders cohort. Collection of blood samples was conducted with approval from the Human Biomaterial Resource Centre (Birmingham, UK), the South-East Scotland Research Ethics Committee (16/SS/0172), and the University of Birmingham Local Ethical Review Committee (ERN 12_0079). Adipose tissue sampling was approved by the Research Ethics Committee (NRES 13/NE/0222) and conducted in accordance with the Declaration of Helsinki. All sample collection, processing, storage, and subsequent experimental procedures complied with Human Tissue Authority guidelines under the Human Tissue Act (2004).

### Sample collection, storage, and processing

#### Blood samples

Blood samples were collected from healthy young (n = 9) and aged (n = 12) donors (**Supp. Table 3**) in 6 ml BD vacutainers containing Lithium Heparin (Scientific Laboratory Supplies, cat. no.: VS367885), Ethylenediaminetetraacetic acid (EDTA) (Scientific Laboratory Supplies, cat. no.: VS367873), silica (Scientific Laboratory Supplies, cat. no.: VS367837), or 3 ml Tempus^TM^ blood tubes (Thermo Fisher Scientific, cat. no.: 4342792).

### Isolation of Serum and Plasma from whole blood

For serum isolation, blood collected in silica BD vacutainers (containing no anti-coagulant) (Scientific Laboratory Supplies, cat. no.: VS367837) was left for 30 min at room temperature (RT) to allow for coagulation before centrifugation at 2000 x g for 10 minutes at RT. The serum layer was aliquoted (500 µl) into cryovials (Greiner, cat. no.: G122263) and stored at -80 °C until further analysis. For plasma isolation, blood collected in EDTA tubes (Scientific Laboratory Supplies, cat. no.: VS367873) was centrifuged at 1800 x g for 8 minutes at RT. The plasma layer was then then transferred to cryovials (Greiner, cat. no.: G122263) in 500 µl aliquots and stored at -80 °C until further analysis.

### Isolation of PBMCs and T cells from whole blood

Peripheral blood mononuclear cells (PBMCs) were isolated by density centrifugation using Ficoll-Paque™ PLUS (Merck, cat. no.: GE17-1440-02). Whole blood was diluted in RPMI media (Fisher Scientific, cat. no.: 11530586) in an 8:1 ratio and layered onto Ficoll-Paque™ PLUS solution in a 2:1 ratio. After centrifugation (400 x g, 30 minutes, RT, no acceleration or break), the PBMC layer was collected from between the Ficoll-Paque™ and plasma layers. The cells were washed using 10 ml of RPMI media (300 x g, 10 minutes, RT) and resuspended in 10 ml of RPMI media once more. The cell counts were obtained by aliquoting a 150 µl of cell suspension in a fresh tube and using a haematology analyser (Sysmex, cat. no.: XN-1000). Using these cell counts, PBMCs designated for later use were separated in a fresh tube, spun again as above, and resuspended at 3 x 10^6^ cells/ml in freezing medium consisting of 10% DMSO (Sigma Aldrich, cat. no.: D2650) in heat-inactivated foetal calf serum (LabTech, cat. no.: FCS-SA/500). The cell suspension was aliquoted at 1 ml in cryovials (Greiner, cat. no.: G122263) which were placed at -80°C storage in a Corning® CoolCell® (Scientific Laboratory Supplies, cat. no.: 432004) overnight and subsequently kept in a storage box at -80°C. PBMCs to be used immediately were kept in RPMI ready for further analysis.

T cells were isolated from PBMCs using the EasySep™ Human T Cell Isolation Kit (Stemcell Technologies, cat. no.: 17951) following the manufacturer’s instructions. In short, PBMCs were resuspended at 5×10^7^cells/ml in MACS buffer (Miltenyi Biotec, cat. no.: 130-091-221) and combined with the human T cell isolation cocktail at 50 µl/ml of cell suspension. After a 5-minute incubation at RT, magnetic dextran RapidSpheres ™ were added at 40 µl/ml, and the mixture was toped up to 5 ml using MACS buffer. The mixture was place in the EasySep™ Magnet (Miltenyi Biotec, cat. no.: 18000) for 3 minutes. The supernatant was poured off into a fresh tube and spun at 300 x g for 10 minutes at RT to obtain the T cell pellet. The T cells were then resuspended in RPMI media (Fisher Scientific, cat. no.: 11530586) with added L-glutamine (Sigma Aldrich, cat. no.: G7513-20ML) and FCS (LabTech, cat. no.: FCS-SA/500) at 5 x 10^5^cells/ml and incubated at 37°C overnight before further analysis.

### Isolation of DNA from whole blood

DNA was isolated from 100 µl of EDTA or heparin blood (fresh or frozen) using a commercially available kit (DNeasy Blood & Tissue Kit, QIAGEN, cat. no.: 69504) following manufacturer’s instructions. Briefly, blood samples were mixed with 20 µl of Proteinase K, 100 µl of PBS (Sigma Aldrich, cat. no.: P4417-50TAB), and 4 µl RNase A (QIAGEN, cat. no.: 19101) for two-minutes at RT. Buffer AL (200 µl) was added and the mixture was incubated at 56°C for 10 minutes before ethanol (200 µl) was added and the sample was transferred into a DNeasy Mini spin column. DNA was captured in the membrane of a spin column by centrifuging at 6000 x g for 1 minute, before washing the membrane with Buffer AW1 (6000 x g for 1 minute) and Buffer AW2 (20000 x g for 3 minutes). The DNA was eluted by addition of 100 µl of Buffer AE and centrifugation at 6000 x g for 1 minute. The elution step was repeated using the eluted sample instead of Buffer AE.

The concentration of the obtained DNA was measured using the Quant-iT™ PicoGreen™ kit (Invitrogen, cat. no.: P7589). DNA samples were measured in duplicates by combining 1 µl of sample with 99 µl of TE buffer and 100 µl of PicoGreen™ working solution in a 96-well clear-bottom plate. Standards (from 0 ng/ml to 1 µg/ml) were created using a serial dilution of DNA stock provided in the kit. Both the samples and standards were measured using a microplate reader (BMG LABTECH, CLARIOstar Plus) with 480 nm excitation and 520 nm emission. Isolated DNA samples were stored at -80 °C until further analysis.

### Isolation of RNA from whole blood

To isolate and validate RNA isolation from blood, blood was collected in EDTA (Scientific Laboratory Supplies, cat. no.: VS367873) and Tempus^TM^ blood tubes (Thermo Fisher Scientific, cat. no.: 4342792). EDTA blood was isolated using QIAamp RNA Blood Mini Kit (QIAGEN, cat. no.: 52304) according to the manufacturer’s instructions. Briefly, blood was mixed with Buffer EL (1:5 ratio), incubated for 15 minutes on ice and centrifuged at 400 x g for 10 minutes at 4 °C. The pellet was washed in Buffer EL (400 x g for 10 minutes at 4 °C) and resuspended in Buffer RLT. Cells were lysed by centrifugation at maximum speed for 2 minutes in the QIAshredder spin column. The generated lysate was mixed with 1 x volume of 70% ethanol and the RNA from this mixture was captured using the QIAamp spin column by spinning for 15 seconds at 8000 x g. The membrane was then washed with 500 µl of Buffer AW1 (15 seconds at 8000 x g), 500 µl of Buffer RPE (15 seconds at 8000 x g), and 500 µl of Buffer RPE (full speed for 3 minutes). Finally, 50 µl of RNase-free water was added to the QIAamp membrane and RNA was eluted by centrifuging for 1 minute at 8000 x g. Samples were stored at -80°C until analysis.

Tempus tubes were stored at RT, 4 °C, or -80 °C for 5, 7, and >7 days respectively (**Supp. Table 4**). RNA was isolated using the Tempus™ Spin RNA Isolation Kit (Thermo Fisher Scientific, cat. no.: 4380204). Tempus™ Blood RNA Tubes were brought up to RT, diluted with 3 ml of PBS (Ca2+/Mg2+-free), vortexed for 30 seconds, and spun at 4 °C (3000 x g for 30 minutes). The supernatant was removed, and RNA pellet was resuspended in 400 µl of RNA Purification Resuspension Solution. RNA Purification Wash Solution 1 was used to pre-wet the membrane of an RNA purification column, and the RNA was captured onto the membrane 16000 x g for 30 seconds at RT. The membrane was washed with RNA Purification Wash Solution 1 (30 seconds at 16000 x g) and two times with RNA Purification Wash Solution 2 (30 seconds at 16000 x g). The membrane was dried by centrifugation for 30 seconds at 16000 x g before RNA was eluted by adding Nucleic Acid Purification Elution Solution, incubating at 70 °C for 2 minutes and centrifugation for 30 seconds at 16000 x g. The elution step was repeated using the collected RNA solution. RNA was transferred into a new collection tube, ensuring the debris pellet is not disturbed and was stored at -80 °C until analysis. Analysis of RNA quantity and quality was performed using the RNA ScreenTape® assay (Agilent Technologies, Santa Clara, CA, USA; cat. No.: 5067-5576) on the Agilent 4200 TapeStation system (Agilent Technologies), and data were processed using TapeStation Analysis Software v5.1.

### Tissue samples

Subcutaneous adipose tissue (SAT) biopsies were obtained from (n = 10) individuals undergoing elective hip-replacement surgery at the Royal Orthopaedic Hospital, Birmingham, UK. Diabetic patients and those taking anti-inflammatory medication within 2 weeks prior to surgery were excluded from the study. Informed consent was obtained from all patients prior to sample collection. Tissue samples were cryopreserved by snap freezing in liquid nitrogen or processed and analysed within less than 5 hours from the time of sample collection. Cryopreserved samples were stored at -80 °C until further analysis.

### Stem cell isolation

Human SAT samples (200 mg) were minced with a scalpel in a culture dish, under a laminar flow hood. Next, samples were transferred to a 50 ml centrifuge tube containing 5 ml of pre-warmed 2 mg/ml Collagenase Type IA (Sigma Aldrich, cat. no.: C9891-1G) prepared in PBS solution (Sigma Aldrich, cat. no.: P4417-50TAB). Tubes were then incubated in a 37 °C in a water bath for 20 minutes, shaking vigorously every 5 minutes. Collagenase digestion was stopped by the addition of 10 mL DMEM (Fisher Scientific, cat. no.: 11584486) supplemented with 10% FCS (LabTech, cat. no.: FCS-SA/500). The digest was pelleted by centrifugation at 800 x g for 10 minutes at RT. Post centrifugation, the upper lipid and oil fractions were carefully removed, and the stromal cell pellet resuspended in pre-warmed DMEM medium supplemented with 20% FCS, 1% L-glutamine (Sigma Aldrich, cat. no.: G7513-20ML), and 1% Pen/Strep solution (Merck, cat. no.: P0781-20ML). Residual lipids were removed by filtering through a 70 µm cell strainer (Corning, cat. no.: 352350) before cells were plated in 2 ml of the supplemented DMEM media in a single well of a 24-well plate (Fisher Scientific, cat. no.: 10380932). The plate was incubated 37 °C (5% CO_2_) until cells reached confluency, with media changed every 2 days.

### Functional assays assessing the hallmarks of ageing

#### Spectral flow cytometry

Frozen PBMCs were thawed in a 37°C water bath for 1 minute, transferred into a fresh 15 ml centrifuge tube, and washed in 10 ml of RPMI-1640 (Fisher Scientific, cat. no.: 11530586) containing 10% FCS (LabTech, cat. no.: FCS-SA/500) and 1% L-glutamine (Sigma Aldrich, cat. no.: G7513-20ML) at 300 x g for 10 minutes at RT. The pellet was resuspended in 10 ml MACS buffer (Miltenyi Biotec, cat. no.: 130-091-221) and the cells were counted using a haematology analyser (Sysmex, cat. no.: XN-1000). For the identification of cell subsets in PBMCs, 3 x 10^5^ cells are required. This number of cells was aliquoted from the PBMC suspension into a round-bottom 96-well plate and spun again as previously described above to obtain a pellet. The remaining cells can be used for other functional assays to maximise the data output from a single sample. The extracellular markers were stained for 30 minutes on ice in the dark, using a mixture of 1 µl of each antibody except for anti-CCR7 (2 µl), 0.5 µl of LIVE/DEAD viability dye, and 19.5 µl of brilliant stain buffer (BD Biosciences, cat. no.: 563794). All supplier details of these antibodies can be found in **Suppl. Table 1**. Following the incubation, the cells were washed in 200 µl of MACS buffer at 250 x g for 5 minutes at 4°C. The pellet was resuspended in 100 µl of fixation/permeabilization solution from the Foxp3/ transcription factor staining kit (Thermo Fisher Scientific, cat. no.: 00-5523-00) and incubated for 30 minutes on ice in the dark. The cells were then washed in 100 µl of diluted permeabilization buffer at 250 x g for 5 minutes at 4°C and resuspended in an intracellular staining cocktail consisting of 0.2 µl of both anti-human p16 and p21 antibodies (**Suppl. Table 1**), and 49.6 µl of permeabilization buffer. The cells were left in the dark on ice for 30 minutes and then washed twice: first in 200 µl of permeabilization buffer (250 x g for 5 minutes at 4°C) and a second time in 200 µl of MACS buffer using the same centrifugation settings. Finally, the cells were resuspended in 200 µl of MACS buffer and were run on the SONY ID7000 spectral flow cytometer (SONY Biotechnology, ID7000™ Spectral Cell Analyzer). Samples were analysed using the ID7000 software (SONY Biotechnology, ID7000™ Software System). Briefly, PBMCs were gated around, and doublets and dead cells were gated out. Utilising the wide array of markers, samples were gated for immune cells and their subsets (T cells, B cells, Monocytes, Macrophages, Dendritic cells and NK cells) and circulating endothelial progenitors (**Suppl. Fig. 1, Suppl. Table 2**).

To confirm the effectiveness of the P16 and P21 antibodies, P8 BJ-5ta fibroblasts (ATCC, cat.no.: CRL-4001) had senescence induced through irradiation. For this, BJ-5ta fibroblasts were cultured in supplemented DMEM media until 80% confluency. At this point, flasks were irradiated with 5Gy of radiation on the CellRad^TM^ (Precision X-ray Irradiation), cultured for a further three days and then stained using the P16, P21 intracellular staining protocol and run on the flow cytometer.

### IMM-AGE score calculation

From the spectral flow cytometry data, 8 immune cell types (total T cells, naive CD4 T cells, effector memory CD4 and CD8 T cells, EMRA CD8 T cells, CD28^−^ CD8 T cells, CD57^+^ CD8 T cells and regulatory T cells) were selected to calculate the IMM-AGE metric as previously described^10,11^ using the RStudio software version 4.5.1 (Posit team (2025). RStudio: Integrated Development Environment for R. Posit Software, PBC, Boston, MA. URL http://www.posit.co/). Following standardisation of the data through a diffusion map, trajectory was built and IMM-AGE scores were calculated based on the location of the cells on the trajectory. The trajectory was dictated by CD28 frequency and scaled between 0 and 1.

### Staining for cellular senescence

For analysis of adipose tissue cellular senescence, SAT biopsies were embedded in cryomolds in OCT (Fisher Scientific, 11912365), frozen on dry ice, and sectioned at 20 µm onto glass slides using a precooled (-30 °C) cryostat (Leica Biosystems, Leica CM1950). The tissue sections were stained for senescence associated β-Galactosidase (SA β-Gal). For this, a hydrophobic pen (Vector Laboratories, cat. no.: H-4000) was used to create a circular border around the tissue section to remove the need to immerse the whole slide into the staining solution and reduce the quantity of reagents used. First, the sections were fixed in a solution of 1% PFA (Sigma Aldrich, cat. no.:HT501128) and 0.2% glutaraldehyde (Thermo Scientific, cat. no.: A17876) diluted in PBS (Sigma Aldrich, cat. no.: P4417-50TAB) for 4 minutes. Then the slide was washed twice with pH 7 PBS and incubated in pH 5.5 PBS for 30 minutes at RT. A β-Gal stain containing 4 mM K_3_Fe(CN)_6_ (Sigma Aldrich, cat. no.: 244023-500G), 4 mM K_4_Fe(CN)_6_ (Sigma Aldrich, cat. no.: P3289-5G), 2 mM MgCl_2_ (Sigma Aldrich, cat. no.: M8266-1KG), and 400 µg/mL X-Gal (C14H15BrClNO6; Thermo Scientific, cat. no.: R0404) dissolved in DMF (Merck, cat. no.: D4551-250ML) was prepared in pH 5.5 PBS and added to the sections. Sections were incubated overnight in the dark at 37 °C without CO_2_. Slides were then fixed with 4% PFA, washed 3 times with pH 7 PBS. Counterstain solution was prepared by combining 1 g of Nuclear Fast Red powder dye (Sigma Aldrich, cat. no.: 60700-5G) and 50 g of potassium aluminium sulphate (Thermo Scientific, A10906) to 500 ml of dH_2_O and brought to boiling temperature. This solution was left to cool at RT overnight and then filtered before application onto the slides using a syringe and 0.22 µm filter (Gilson, cat. no.: ANP2522). Finally, coverslips were mounted using Immuno-Mount^TM^ solution (Fisher Scientific, cat. no.: 10662815) and a slide scanner (Zeiss, Axio Slide Scanner 7) microscope was used to image the slides. Analysis was performed using FIJI software^64^, calculating the percentage of β-Gal-stained cells compared to total cells within the section. Whole tissue staining was also performed by taking 50 mg of tissue and incubating overnight submerged in the pH 5.5 β-Gal solution described above in a dark at 37 °C without CO_2_. Tissue was then fixed in 4% PFA for 30 minutes and imaged. Images were analysed using FIJI by isolating the cyan image and intensity of cyan pixels were determined by drawing a region of interest using FIJI’s standard measurement functions.

### Quantification of serum and plasma cytokine concentrations

Serum and EDTA plasma concentration of CXCL-9, GDF-15, IL-6 and TNFα, were measured using commercially available ELISA kits (CXCL9; DY392-05, GDF15; DY957, IL-6; DY206, TNF; DY210, R&D Systems, MN, USA) following the manufacturer’s protocol. Firstly, capture antibody was diluted in PBS (pH 7.2) as specified by the manufacturer and added to the required wells of an uncoated nunc-ImmunoT ELISA plate (cat.no. M9410-1CS, ThermoFisher Scientific, MA, USA) overnight at RT. The following day, wells were washed 3 times with 300 µl wash buffer (PBS pH 7.2, containing 0.05% Tween® 20 cat.no: P9416, Sigma, Uk) and then blocked with reagent diluent (PBS containing 1% BSA cat.no: 9418, Sigma, UK) for 1 hour. Plates were then aspirated and washed as described above. Next, protein standards were serially diluted 2-fold in reagent diluent. Standards and samples were then added to the plate in duplicate and incubated for 2 hours at RT. Following incubation, plates were again washed as above, before the addition of capture antibody (diluted in reagent diluent to manufactures specification) for 2 hours at RT. Streptavidin-HRP (diluted in reagent diluent) was then added to wells for 20 minutes at RT, in the dark. Following a final wash step as above, substrate solution was added to wells for up to 20 minutes. Finally, reactions were stopped by the addition of stop solution (2N H2SO4) and optical density immediately measured at 450 nm using a Varioskan Lux plate reader (ThermoFisher Scientific). Optical imperfections in the plate were limited by correcting with absorbance measured at 570 nm. The average absorbance for each standard was used to generate a standard curve using GraphPad Prism v9 statistical package. Concentrations of unknown samples were then calculated using this standard curve and expressed as pg/ml.

### Measurement of metabolic flux in human T-Cells

The metabolic flux was measured using the Seahorse Analyser (Agilent, Seahorse XFe96 Analyzer). For this, T cells were isolated as described above from PBMCs freshly obtained from heparin or EDTA blood. T cells were incubated overnight a 37°C, 5% CO_2_ with or without CD3/CD28 stimulation (Stemcell technologies, cat. no.: 17951) at 3 million cells per well of a 12 well plate in RPMI media (Fisher Scientific, cat. no.: 11530586) supplemented with 10% FCS (LabTech, cat. no.: FCS-SA/500) and 1% L-glutamine (Sigma Aldrich, cat. no.: G7513-20ML). After the incubation, the cells were counted using a haematology analyser (Sysmex, cat. no.: XN-1000) and pelleted in 1.5 ml centrifuge tubes. Subsequently, the cells were resuspended in 300 µl of Complete assay media (CAM) composed of Seahorse RPMI media (Agilent, cat. no.: 103576-100), 2 mM L-glutamine (Sigma Aldrich, cat. no.: G7513-20ML), 1 mM sodium pyruvate (Sigma Aldrich, cat. no.: P8574-5G), and 10 mM glucose (Sigma Aldrich, cat. no.: G8270-100G). From this suspension, 200000 cells were plated onto the wells of the Seahorse assay cell culture plate (Agilent, cat. no.: 103723-100) that was previously coated for 30 minutes with a 50 µg/ml poly-D-lysine solution (Scientific Laboratory Supplies, cat. no.: A-003-E). The plate was spun at 250 x g for 2 minutes (no brake), the wells were topped up to 180 µl of CAM, and the plate was incubated at 37°C, 5% CO_2_ for 1 h. The Seahorse cartridge plate (Agilent, cat. no.: 103723-100) was prepared the day before by incubating overnight with the Seahorse calibrant solution (200 µl/well) (Agilent, cat. no.: 103723-100). After the overnight incubation, the cartridge plate was prepared for the assay by adding 20 µl of 20 µM oligomycin (Merck, cat. no.: O4876-25MG) into port A, 22 µl of 30 µM BAM15 (Merck, cat. no.: SML1760-5MG) into port B, 25 µl of 10 µM rotenone (Merck, cat. no.: R8875-5G) + antimycin A (Merck, cat. no.: A8674-100MG) into port C, and 27 µl of 200 µM monensin (Merck, cat. no.: M5273-500MG) into port D. The Seahorse assay was set up as follows: 4 cycles of 3 minutes of mixing and 3 minutes of measuring for the baseline oxygen consumption rate (OCR) and extra-cellular acidification rate (ECAR) levels, followed by 3 cycles of 3 minutes of mixing and 3 minutes of measuring for each individual treatment in the order listed above. The results were normalised by visualising each well under a brightfield microscope and the cell numbers were quantified using the particle count function from the FIJI software^64^. The cell counts were averaged for all wells, and the averaged number was then used to divide each individual count to obtain the normalisation factor for each well. The data was then processed using the Seahorse analytics software (Seahorse Analytics Software, version 1.0.0-768, Agilent Technologies, G3299-90000) and the final outputs were calculated from the normalised results using the following equations:

| **Basal glycolysis** | **the average of the baseline ECAR readouts** |
| --- | --- |
| **Maximal glycolytic capacity** | the average of the readouts after monensin |
| **Compensatory glycolysis** | maximal glycolytic capacity - basal glycolysis |
| **ATP restricted glycolysis** | maximal glycolytic capacity - compensatory glycolysis |
| **ATP supply rate** | mitoATP supply rate + glucoATP supply rate |
| **mitoATP supply rate** | $\text{OCR ATP} \times 2 \times 2.75$ (P/O for mixed substrates) |
| **glycoATP supply rate (=glycoPER)** | PER - mitoPER |
| **PER** | baseline ECAR * 2.8 (Buffer factor) * volume of well * 1.6 (Kvol for XF96) |
| **mitoPER** | mitoOCR * 0.61 (CCF for XF96) |
| **Basal mitochondrial respiration** | baseline - R/A |
| **Phosphorylation-linked respiration** | basal mito respiration - (oligomycin-R/A) |
| **Proton leak** | oligomycin - R/A |
| **Coupling efficiency** | $(1 - (\text{proton leak} / \text{Basal mito respiration})) \times 100$ |
| **Maximal respiration** | BAM15 - R/A |
| **Spare capacity** | $((\text{Max respiration} / \text{basal mito respiration}) - 1) \times 100$ |
| **Non-mitochondrial respiration** | R/A read out |
| **ATP-linked respiration** | Basal mito respiration - oligomycin |
| **mitoOCR** | basal mito respiration - R/A |
| **OCR ATP** | baseline OCR - oligomycin |

### Measurement of metabolic dependencies in human PBMCs

The SCENITH method^33^ was adapted and optimised to measure metabolic dependencies. For this, frozen PBMCs were thawed and washed as described. Cells were then resuspended in RPMI media (Fisher Scientific, cat. no.: 11530586) supplemented with 10% FCS (LabTech, cat. no.: FCS-SA/500) and 1% L-glutamine (Sigma Aldrich, cat. no.: G7513-20ML) at 200,000 cells in 100 µl per tube (Fisher Scientific, cat. no.: 10579511). Separate tubes were prepared per sample – 7 tubes to be treated with different inhibitors, 1 tube for untreated control, 1 tube for unstained cells (autofluorescence control), and 1 tube for isotype control. The inhibitor solutions were prepared as follows: 1 M solution of 2DG (Merck, cat. no.: 25972-1GM) in PBS (Sigma Aldrich, cat. no.: P4417-50TAB); 6320 µM Oligomycin (Merck, cat. no.: O4876-5MG) stock in DMSO (Merck, cat. no.: D8418-50ML) - further diluted to 500 µM in PBS for working solution; 9529 µM BPTES (Scientific Laboratory Supplies, cat. no.: SML0601-5MG) stock in DMSO - further diluted to 1000 µM PBS for working solution; 14759 µM ETO (Cambridge BioScience, cat. no.: 11969-10mg-CAY) stock in PBS - further diluted to 500 µM in PBS for working solution; and 5 mg/ml Harringtonine (Cambridge BioScience, cat. no.: CAY15361-5 mg) stock in DMSO - further diluted to 2 mg/ml in DMSO for working solution.

The cells were then treated as with a final concentration of 100 mM of 2DG, 5 µM of oligomycin,10 µM of BPTES, 5 µM of ETO, and 2 µg/ml of Harringtonine in individual tubes. A tube with vehicle control (1:100 DMSO) and a tube combining all the inhibitors were also prepared, as well as five untreated tubes for compensation measurement. All of these were incubated with the respective treatments for 30 minutes at 37 °C, 5% CO_2_. Subsequently, a puromycin (Merck, cat. no.: P7255-25MG) treatment (10 µg/ml) was applied for all tubes for 30 minutes at 37 °C, 5% CO_2_. The cells were then washed twice in 200 µl of PBS (250 x g for 5 minutes at 4 °C). Staining solution (per tube) was prepared using 1 µl of CD3 PECy7 (Invitrogen, cat. no.: 25-0038-42, clone: UCHT1), CD4 BV421 (BioLegend, cat. no.: 300532, clone: RPA-T4), CD45RA APC (BioLegend, cat. no.: 304112, clone: HI100) antibodies, 2 µl of CCR7 FITC (BioLegend, cat. no.: 353216, clone: G043H7) antibody, and 40 µl of MACS buffer (Miltenyi Biotec, cat. no.: 130-091-221). Individual antibodies and up to 50 µl of MACS buffer were used to stain the extracellular compensation controls, and 50 µl of MACS was used for the unstained controls, intracellular compensation, and the isotype control. After an incubation on ice in dark for 20 minutes, the cells were washed as before (250 x g for 5 minutes at 4 °C) and resuspended in a mixture of 100 µl of PBS and 150 µl of Cyto-Fast Fix/Perm buffer from the Foxp3/ transcription factor staining kit (Thermo Fisher Scientific, cat. no.: 00-5523-00). The cells were permeabilised at RT in dark for 20 minutes and after the addition of 500 µl of Cyto-Fast Perm wash solution, spun at 250 x g for 5 min at 4 °C. A cocktail of 92 µl of Perm wash solution and 5 µl of Tru-Stain (BioLegend, cat. no.: 422302) was applied for 10 minutes in the dark at RT and subsequently, 3 µl of anti-puromycin AF488 antibody (BioLegend, cat. no.: 381506, clone: 2A4) was added to each tube except for the unstained control, extracellular compensation, and isotype control. The isotype control tube was stained using 3 µl of IgG2a isotype control antibody (BioLegend, cat. no.: 400233, clone: MOPC-173). All tubes were kept on ice in the dark for 1 hour and were subsequently washed in 200 µl of MACS (250 x g for 5 minutes at 4 °C). Post centrifugation, cells were resuspended in 300 µl of MACS buffer and flow cytometry was performed using the LSR Fortessa analyser (BD Biosciences, LSRFortessa™ X-20 Cell Analyzer). Using FlowJo software (BD Biosciences, FlowJo™ v11 Software), PBMCs were gated for single cells, and subsequently for CD3^+^CD4^+^ T cells and key subsets - naive, central memory (CM), effector memory (EM), and effector memory CD45RA re-expressing cells. For all subsets metabolic dependencies were calculated from the median expression of puromycin within different inhibited tubes using the following equations: Glucose dependence = 100x((veh-2DG)/(veh-AIs)). Mitochondrial dependence = 100x((veh-oligo)/(veh-AIs)). Glycolytic capacity = 100-(100x((veh-oligo)/(veh-AIs))). FAO+AAO capacity = 100-(100x((veh-2DG)/(veh-AIs))). Abb: Veh; vehicle control, AIs; all inhibitors, oligo; oligomycin, FAO; fatty acid oxidation, AAO; amino acid oxidation.

### DNA copy number

The relative DNA copy number was estimated through a qPCR assay^33^. The reactions were performed using 5 µl of the PowerUp™ SYBR™ Green Master Mix (Thermo Fisher, cat. no.: A25742), 1 µl of forward primer (10 µM solution), 1 µl of reverse primer (10 µM solution), 2 µl of DNA template (6 ng/µl dilution), and 1 µl of nuclease-free water (Thermo Fisher, cat. no.: AM9920). Primers were obtained from Merck custom DNA Oligo service (cat. no.:41105327) using the sequences presented in **Supp. Table 5**. The real time PCR system CFX Opus 348 (Bio-Rad, Hertfordshire, UK) was set up as follows: 2 min at 50 °C, 2 min at 95 °C, followed by 40 cycles of: i. 15 sec at 95 °C, ii. 30 sec at 61 °C with fluorescence recording, melt curve from 60-95 °C. The assay was run in triplicates for all genes and samples. If Ct value of any of the replicates differed from the other two by 0.5 or more, the outlier was removed from the analysis. To obtain the relative mtDNA copy number, the Ct values for the nuclear genes from all samples were averaged together. This total average was then used to create the normalisation factor by dividing the individual Ct averages of nuclear genes of each sample. The averages the Ct of the mitochondrial genes for each individual sample were then divided by their respective normalisation factor and the relative mtDNA copy number was obtained.

### Assessment of mTOR signalling in human PBMCs

PBMCs isolated and cryopreserved as described above. The cells were thawed in a 37 °C water bath for 1 minute and washed in 10 ml of RPMI-1640 (Fisher Scientific, cat. no.: 11530586) containing 10% FCS (LabTech, cat. no.: FCS-SA/500) and 1% L-glutamine (Sigma Aldrich, cat. no.: G7513-20ML) at 300 x g for 10 minutes at RT. The pellets were resuspended up to 1 million cells / ml in pre-warmed supplemented RPMI and 200 µl of this suspension was plated into a 96-well round bottom plates and incubated for 24 h at 37 °C, 5% CO_2_. The PBMCs were then stimulated with 0.5 µg/ml PMA (Merck, cat. no.: P1585-1MG) and 0.5 µg/ml Ionomycin (Merck, cat. no.: I0634-1MG) for 30 minutes at 37 °C, 5% CO_2_. Next, the cells were pelleted at 250 x g for 5 minutes and then resuspended in 50 µl of extracellular antibody cocktail consisting of 1 µl of anti-human CD3 PeCy7 (Invitrogen, cat. no.: 25-0038-42, clone: UCHT1),1 µl of anti-human CD4 BV421 (BioLegend, cat. no.: 300532, clone: RPA-T4), 1 µl of anti-human CD45RA APC (BioLegend, cat. no.: 304112, clone: HI100), 2 µl of anti-human CCR7 FITC (BioLegend, cat. no.: 353216, clone: G043H7), and 45 µl of MACS buffer (Miltenyi Biotec, cat. no.: 130-091-221). The cells were incubated with for 20 minutes in the dark, on ice. Post incubation, wells were washed with 200 µl of MACS buffer for 5 minutes at 250 x g at 4 °C and fixed by the addition of 100 µl of fix/perm solution from the Foxp3/ transcription factor staining kit (Fisher Scientific, cat. no.: 00-5523-00) for 30 minutes in the dark, on ice. Cells were then washed with 100 µl of 1X permeabilization buffer (5 minutes, 250 x g, at 4 °C) and resuspended in 50 µl of permeabilization buffer with 1 µl of PE Phospho-S6 antibody (Fisher Scientific, cat. no.: 12-9007-42, clone: cupk43k) for 20 min on ice in the dark. Cells were then washed with 250 µl of 1X permeabilization buffer (250 x g for 5 minutes at 4 °C), followed by a second wash with 250 µl of MACS buffer (250 x g for 5 minutes at 4 °C). Post centrifugation, cells were resuspended in 300 µl of MACS buffer and flow cytometry was performed using the LSR Fortessa analyser (BD Biosciences, LSRFortessa™ X-20 Cell Analyzer), collecting data on at least 20,000 CD4 T cells per sample. Using FlowJo software (BD Biosciences, FlowJo™ v11 Software) PBMCs were gated for single cells, and subsequently for CD3^+^CD4^+^ T cells and key subsets - naive, central memory (CM), effector memory (EM), and effector memory CD45RA re-expressing cells. All subsets were analysed for median expression of Phospho-S6.

### Measurement of autophagic flux in human T Cells

PBMCs were thawed and washed in the same manner as described for the mTOR signalling assay. From these, T cells were isolated using the protocol described above and incubated overnight at 37 °C, 5% CO_2_ in a round bottom 96 well plate (40,000 cells/well). Post incubation, cells were treated with 100 µM chloroquine (Merck, cat. no.: C6628-25G) diluted in cell culture-grade water and incubated for 2 hours at 37 °C and 5% CO_2_. Post treatment, T-cells were pelleted in the plate (250 x g for 5 minutes at 4 °C), permeabilised in 100 µl of 0.1% saponin (Fisher Scientific, cat. no.: 8047-15-2) diluted in MACS buffer (Miltenyi Biotec, cat. no.: 130-091-221), and with 2 µl of one of the anti-LC3II antibodies (CYTEK, Guava® Autophagy kit FITC antibody, cat. no.: CS208214, clone: 4E12; R&D systems, cat. no.: IC9390R, clone: 1251A) for 30 minutes on ice in the dark. T-cells were washed in MACS buffer, removing non-autophagosome incorporated LC3II (250 x *g* for 5 minutes at 4 °C), resuspended in 200 µl of MACS buffer, and transferred to Falcon™ FACS tubes (Fisher Scientific, cat. no.: 10579511). The flow cytometric analysis was performed using SR Fortessa analyser (BD Biosciences, LSRFortessa™ X-20 Cell Analyzer) to measure abundance of LC3II. Samples were analysed in FlowJo (BD Biosciences, FlowJo™ v11 Software) for medium LC3II expression.

### Measured of mRNA expression of DNA damage and repair related genes

One-step quantitative real-rime polymerase chain (qRT-PCR) was performed using 5 ng of RNA (extracted as described above), using SYBR green chemistry in a 384 well PCR plate (BioRad cat.no.: HSP3801). The constituents per reaction are detailed in **Supp. Table 6** and primer sequences are reported in **Supp. Table 5**. All reactions were performed using a CFX Opus Real-Time PCR System (Bio-Rad, Hertfordshire, UK) as described in **Supp. Table 7**. Samples were assayed in triplicate and a non-template control was included for each gene of interest to identify the formation of primer-dimers. Ct values were determined as the point where the curve crossed the defined threshold in the exponential phase of RNA amplification. Melt curves were included for all reactions to confirm the amplification of a single product. The relative expression of the target gene was calculated using the 2^^(-ΔCt)^ method, normalised to the housekeeping gene and relative to the young cohort.

### Stem cell differentiation and characterisation

Differentiation into adipocyte or osteoblast lineages was achieved by replacing alpha-MEM medium with the relevant differentiation media (adipogenic media: cat.no.: PT 4135, Lonza. Osteogenic media: cat.no.: PT 4120, Lonza). Confirmation of osteogenic differentiation was quantified by staining of mineralised nodules using an alizarin red solution (40 mM alizarin red, cat.no.: A5533, Sigma, Gillingham, UK) in 1% ammonia hydroxide at pH 4.5. Following a 25 min incubation at RT, cells were washed with PBS to remove excess stain and imaged using brightfield microscopy. Confirmation of differentiation into mature adipocytes was performed using oil red O staining. A 0.5% oil red O solution was prepared by dilution of oil red O (cat.no. 12989.22, ThermoFisher) in 100% Isopropanol (cat. no.: 190764, Sigma). This Oil Red O solution was diluted in distilled water (3:2) to create a working stain. Oil red O working solution was left to stand at RT for 10 minutes before filtering through a 0.22um filter. Tissue sections, (prepared as above) were fixed with 4% PFA for 5 minutes, and washed twice with H_2_0 (1 minute immersion, RT). Next, tissue sections were immersed in 60% isopropanol for 2 minutes, followed by immediate immersion in oil red O working solution for 15 minutes, at RT. Tissue sections were then rinsed in 60% isopropanol and again washed twice with H_2_O For 3 minutes, Harris haematoxylin (cat.no: HHS128, Sigma, UK) for 1 minute. Finally, tissue sections were washed twice with H₂O for one minute before application of a drop of aqueous mountant and cover slip. Stained sections were imaged using brightfield microscopy immediately after mounting.

### Statistical Analysis

Data were analysed using GraphPad Prism (v9) and R (v4.5.1) and presented as mean ± SEM for n independent experiments. Sample sizes are indicated in the figure legends, and each data point represents one biological replicate unless otherwise stated. In experiments with technical replicates, data points represent the mean of technical replicates derived from a single biological sample. No data were excluded from the analyses. Normality was assessed using Shapiro-Wilk test. Univariate analysis was performed using paired, unpaired t-test, paired t-test or Mann Whitney U test. Multivariate analysis was performed using ANOVA with Šídák’s post-test. p<0.05 was deemed statistically significant.

