## Supplementary Figures and Tables for "Translational toolkit for reproducible, cross-study profiling of human ageing hallmarks in human blood and tissue"

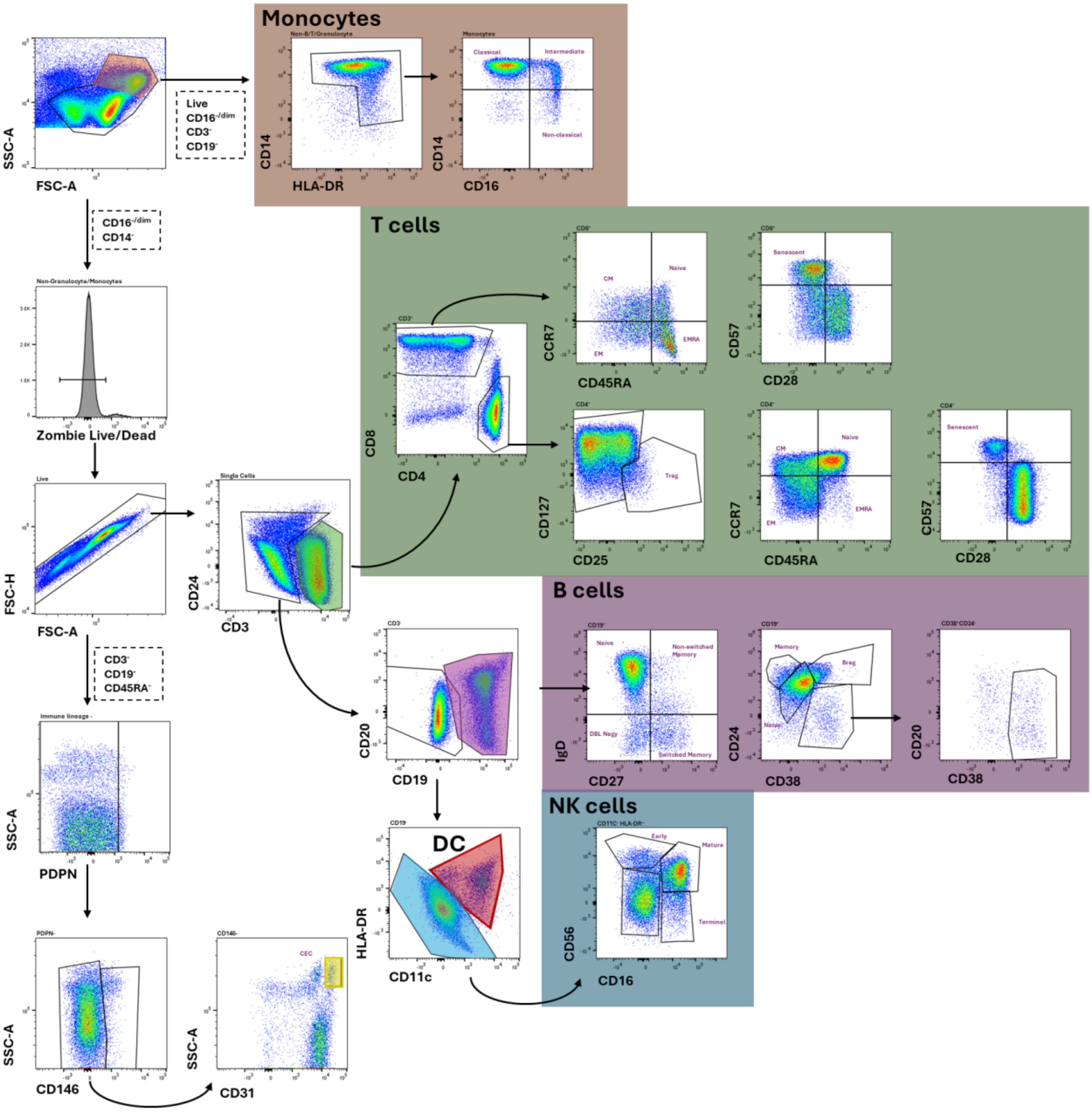


**Supplementary Figure 1. Gating strategy used to define immune cell subsets following spectral flow cytometry.**

Dot plots showing the gating strategy used to identify subsets of monocytes, T cells, B cells, natural killer (NK) cells, dendritic cells (DC), and circulatory endothelial cells (CEC) from PBMCs. The breakdown of all markers used to identify cellular subsets are also presented in Supplementary table 2.

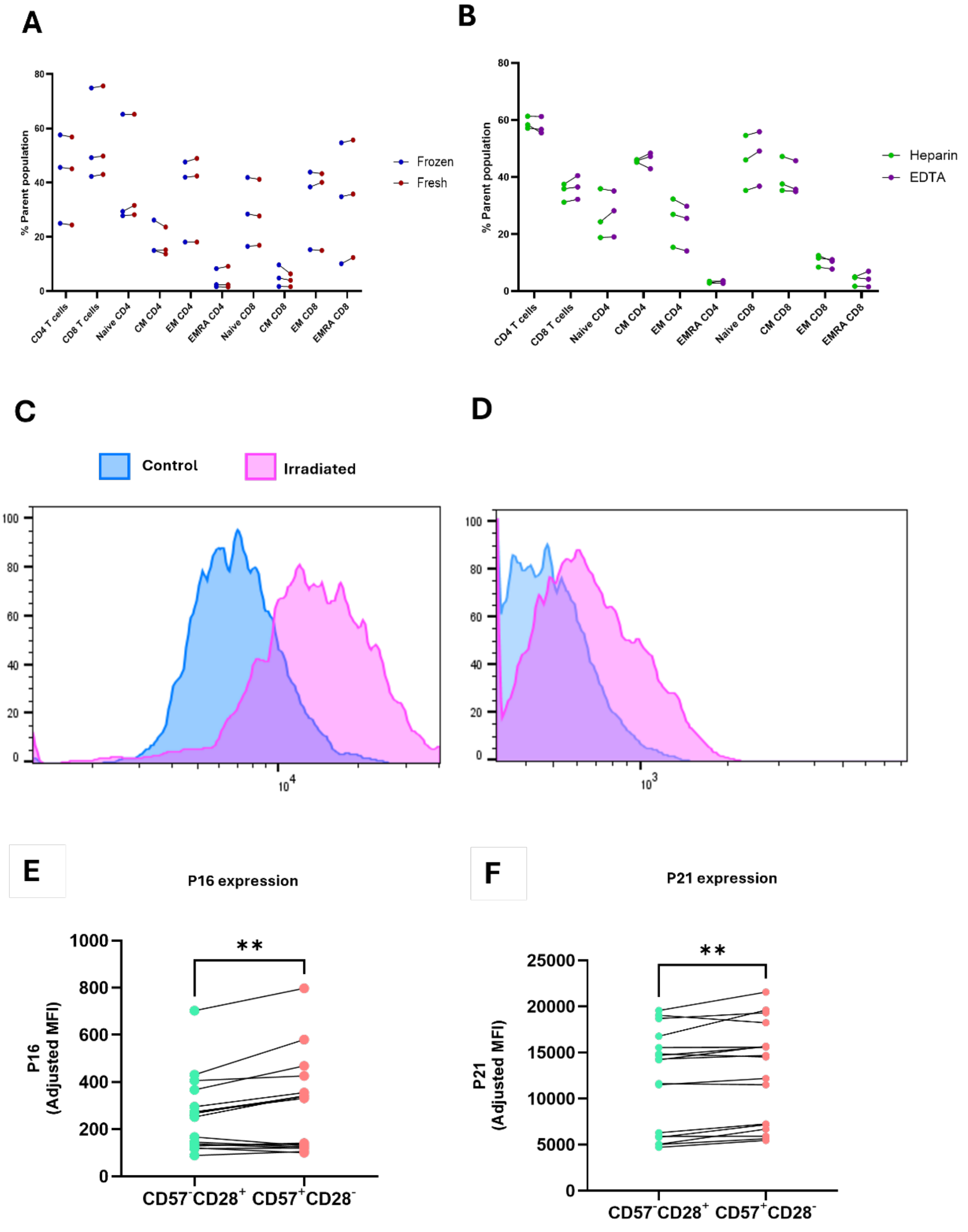


**Supplementary Figure 2. Optimisation and validation spectral flow cytometry panel**
Comparison of T cell subsets identified by spectral flow cytometry in patient matched freshly **(A)** isolated (blue) and cryopreserved (red) PBMCs and from **(B)** frozen cells collected in heparin (green) or EDTA (ethylenediaminetetraacetic acid, purple) anti-coagulant tubes. Representative histograms showing intracellular **(C)** P16 and **(D)** P21 Expression in BJ-5ta fibroblasts at baseline and following irradiation to induce cellular senescence. **(E)** P16 and **(F)** P21 expression as defined by mean fluorescence intensity (MFI) in non-senescent ($CD57^-CD28^+$ - green) and senescent ($CD57^+CD28^-$ - red) T cells from aged participants (n=12). ** denotes p<0.01 by paired t-test.

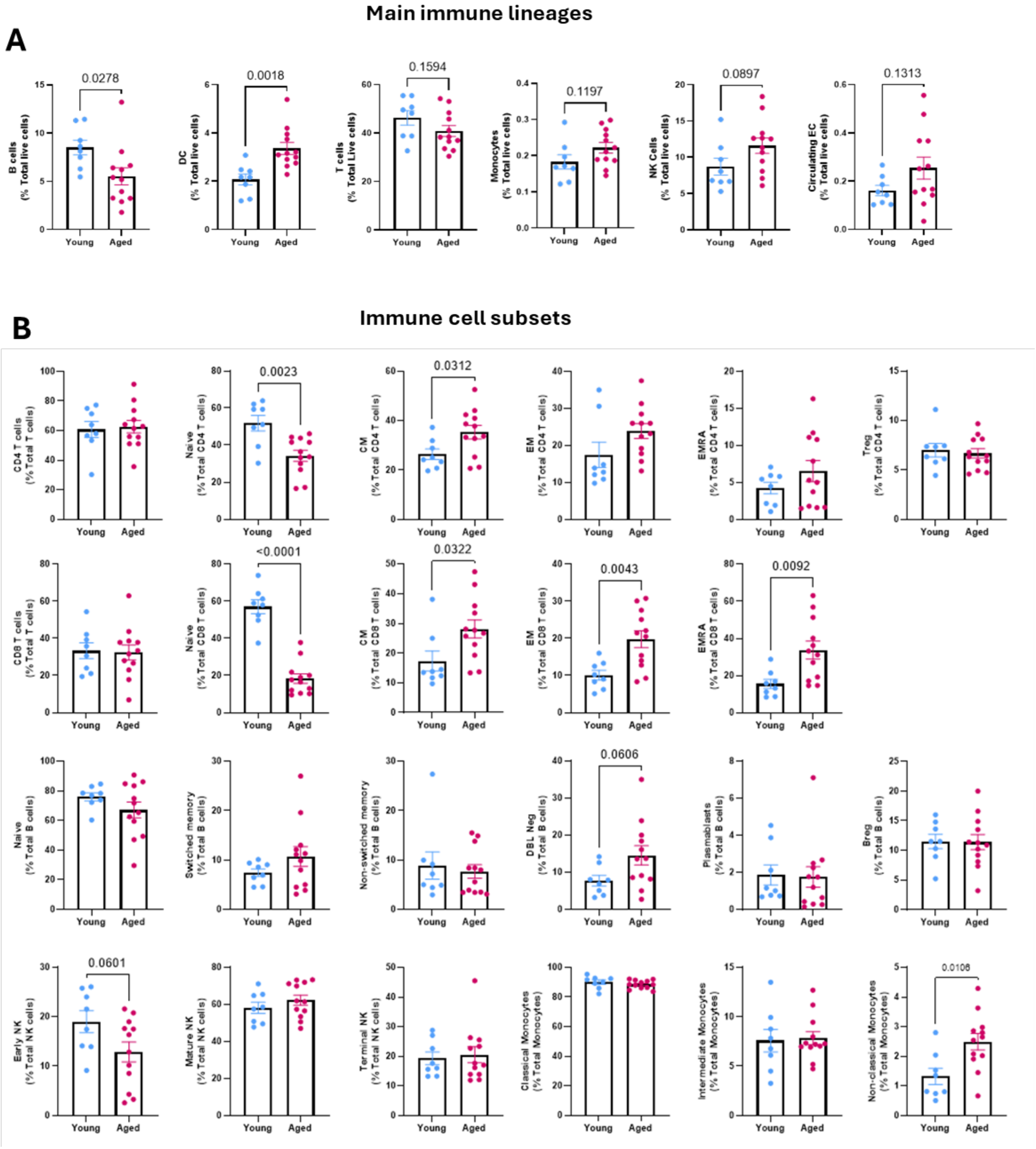


**Supplementary Figure 3. Main lineage and immune cell subsets identified by spectral flow cytometry in young and aged participants.**

**A.** B cells, Dendritic cells, T cells, monocytes, NK cells and circulating endothelial cells in young and aged participants. Cell populations are expressed as % of total live cells. **B.** All CD4+ and CD8+ T cell, B cell, NK and monocyte subsets identified using the gating stratify presented in Supplementary Figure 1 and are presented as % of bulk lineage population. n=8 young and n=12 aged participants. Data are presented as mean ± S.E.M. Statistical significance was assessed in all graphs by unpaired T-tests following normality testing.

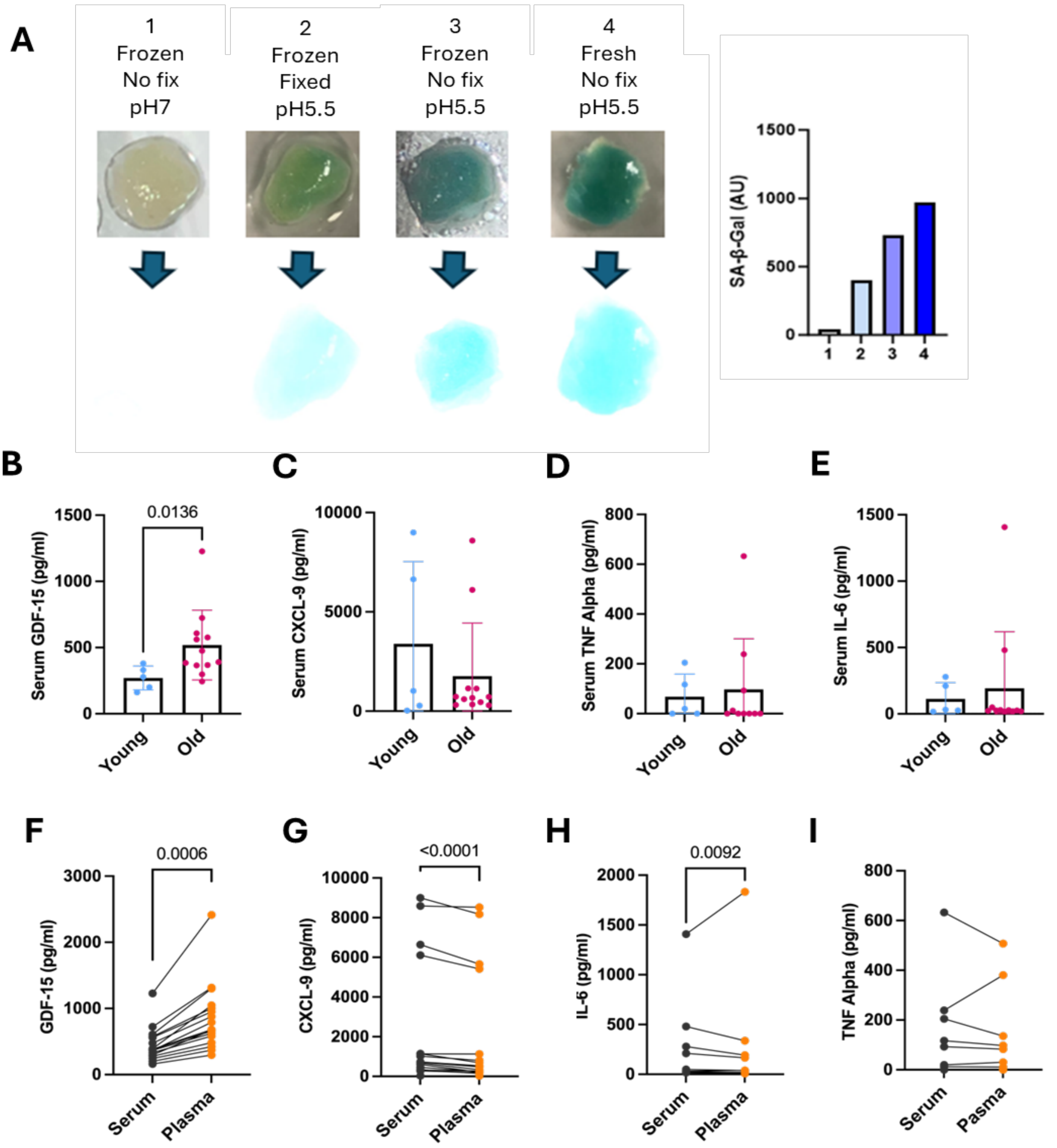


**Supplementary Figure 4. Senescence assessments in tissues and circulating plasma**
**A.** (left) Photographic images of human subcutaneous adipose tissue following SA-beta gal staining under different conditions: (1) subcutaneous adipose tissue processed, fresh at pH7; (2) processed following fixing with paraformaldehyde and cryopreservation at -80; (3) pH5.5; processed following cryopreservation at -80, pH 5.5 and (4) processed Fresh, pH5.5. (Right) Corresponding bar chart of adipose tissue cyan pixel density calculated using Image J, where the x-axis numbers represent the conditions numbered in the images. Concentration of **B.** GDF-15, **C.** CXCL-9, **D.** TNF Alpha and **E.** IL-6 in the serum of young (n=5 and aged (n=12) participants. Concentration of **F.** GDF-15, **G.** CXCL-9, **H.** IL-6 and **I.** TNF alpha in patient matched serum and plasma samples from young (n=5) and aged (n=12) participants. Data are mean ± S.E.M. Statistical significance was assessed (B-E) unpaired Mann Whitney U or (F-I) paired Wilcoxon matched-pairs signed rank test.

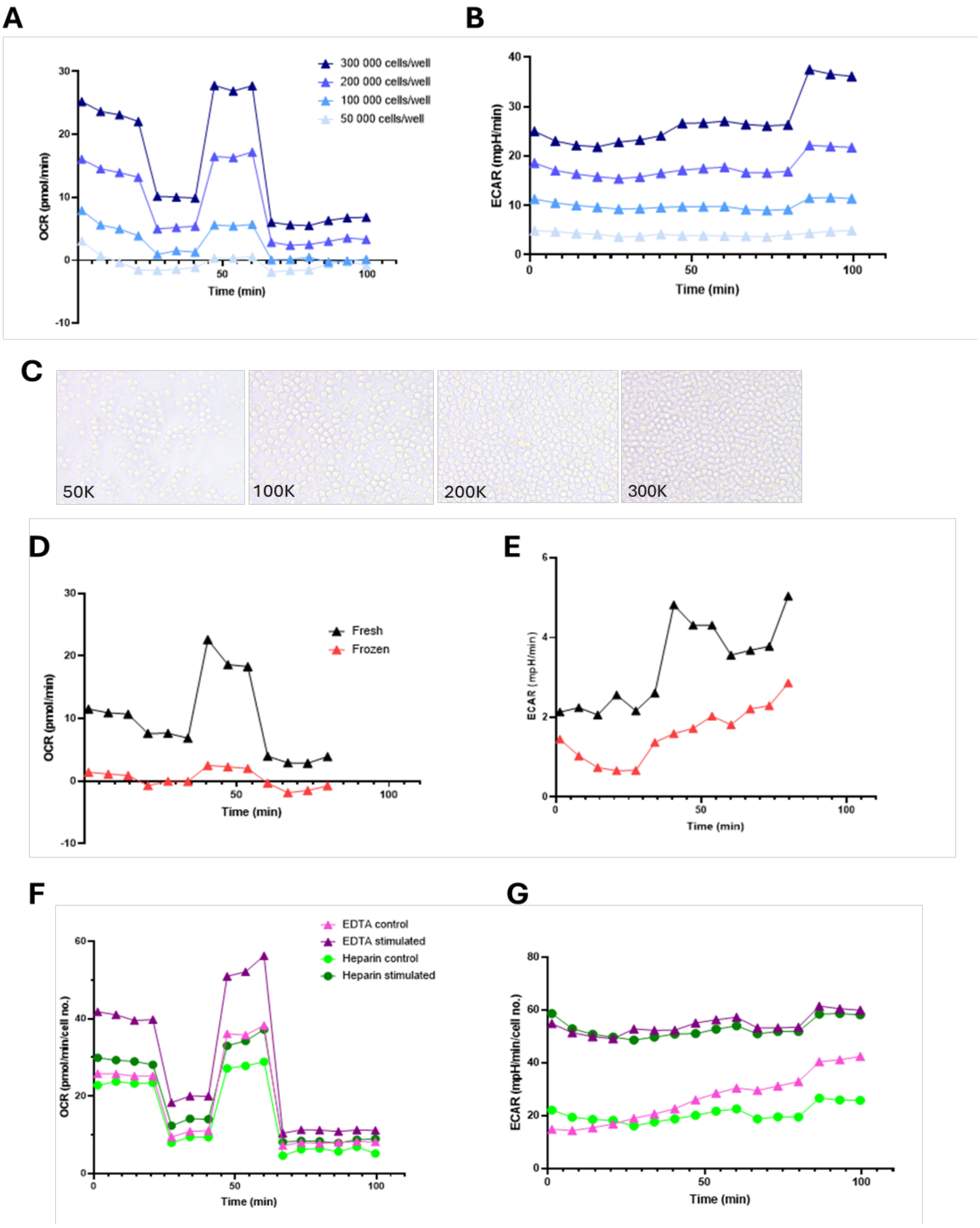


**Supplementary Figure 5. Optimisation of seahorse experimental parameters.**
**A.** OCR and **B.** ECAR traces from titration experiments using T cells derived from young participants (n=3 biological replicates). **C.** Representative well images post completion of seahorse T-cell titration experiments. **D.** OCR and **E.** ECAR traces from freshly isolated and frozen T cells, obtained from a young participant (n=3 biological replicates). **F.** OCR and **G.** ECAR traces from T cells obtained from blood collected with EDTA and heparin blood tubes, with and without 24h stimulation with CD3/CD28 (n=3 biological replicates).

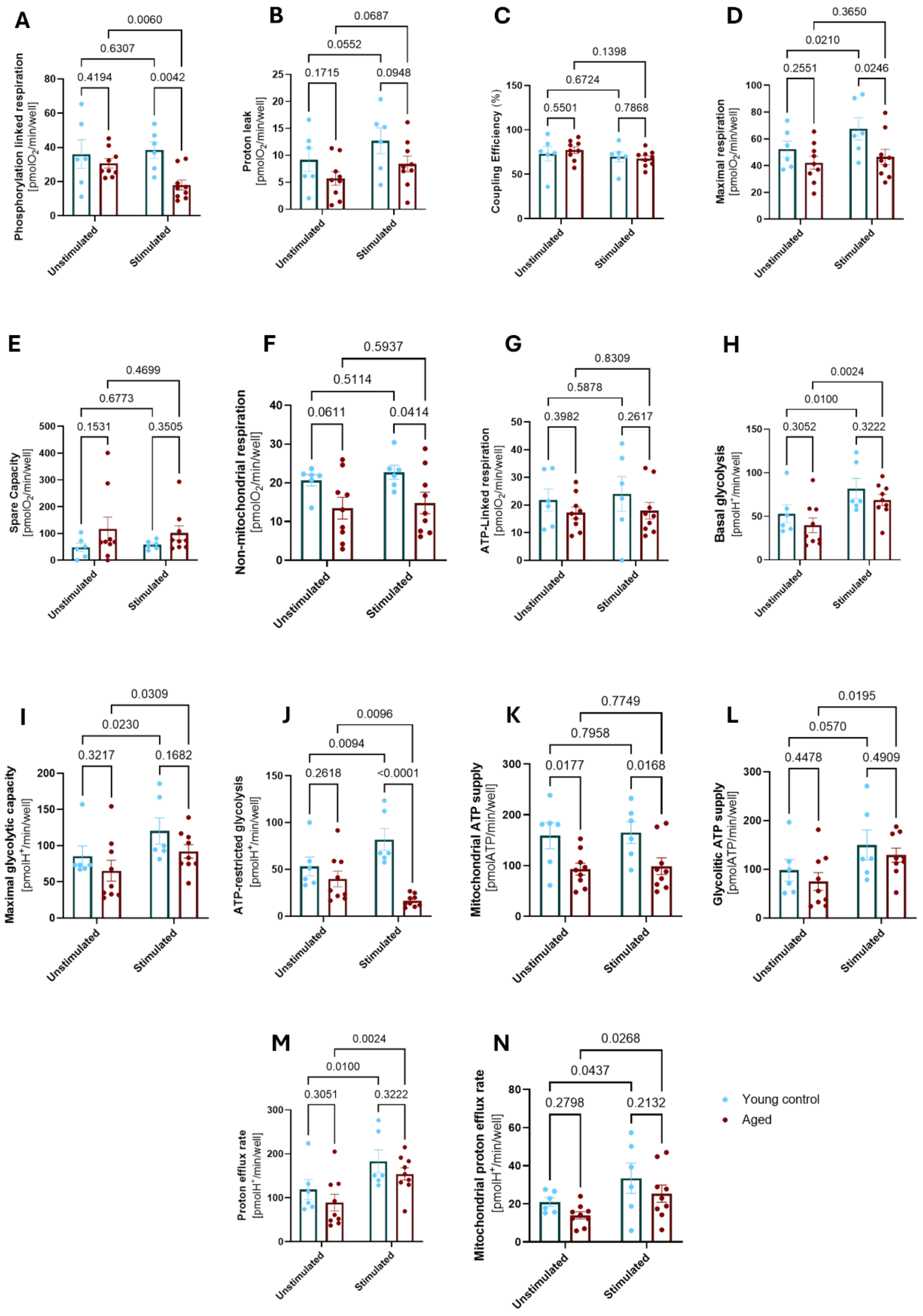


**Supplementary Figure 6. Full metabolic readouts from seahorse assays performed on T-cells from young and aged participants.** T cells were isolated from young (blue) and aged (red) donors and stimulated with or without CD3/CD28 for 24 hours prior to analysis using seahorse. **A.** Phosphorylation Linked respiration, **B.** Proton Leak. **C.** mitochondrial coupling efficiency. **D** Maximal mitochondrial respiration. **E.** Mitochondrial spare capacity. **F.** Non-mitochondrial respiration. **G.** ATP-Linked respiration. **H.** Basal glycolysis. **I** Maximal glycolytic capacity. **J.** ATP-Restricted glycolysis. **K.** Mitochondrial ATP-Production. **L** Glycolytic ATP production. **M.** Proton efflux rate. **N.** Mitochondrial proton efflux rate all expressed as (pmol/min/well). Data are mean ± S.E.M for n=6 young and n=9 aged participants. Statistical significance was determined by Sidak post-test.

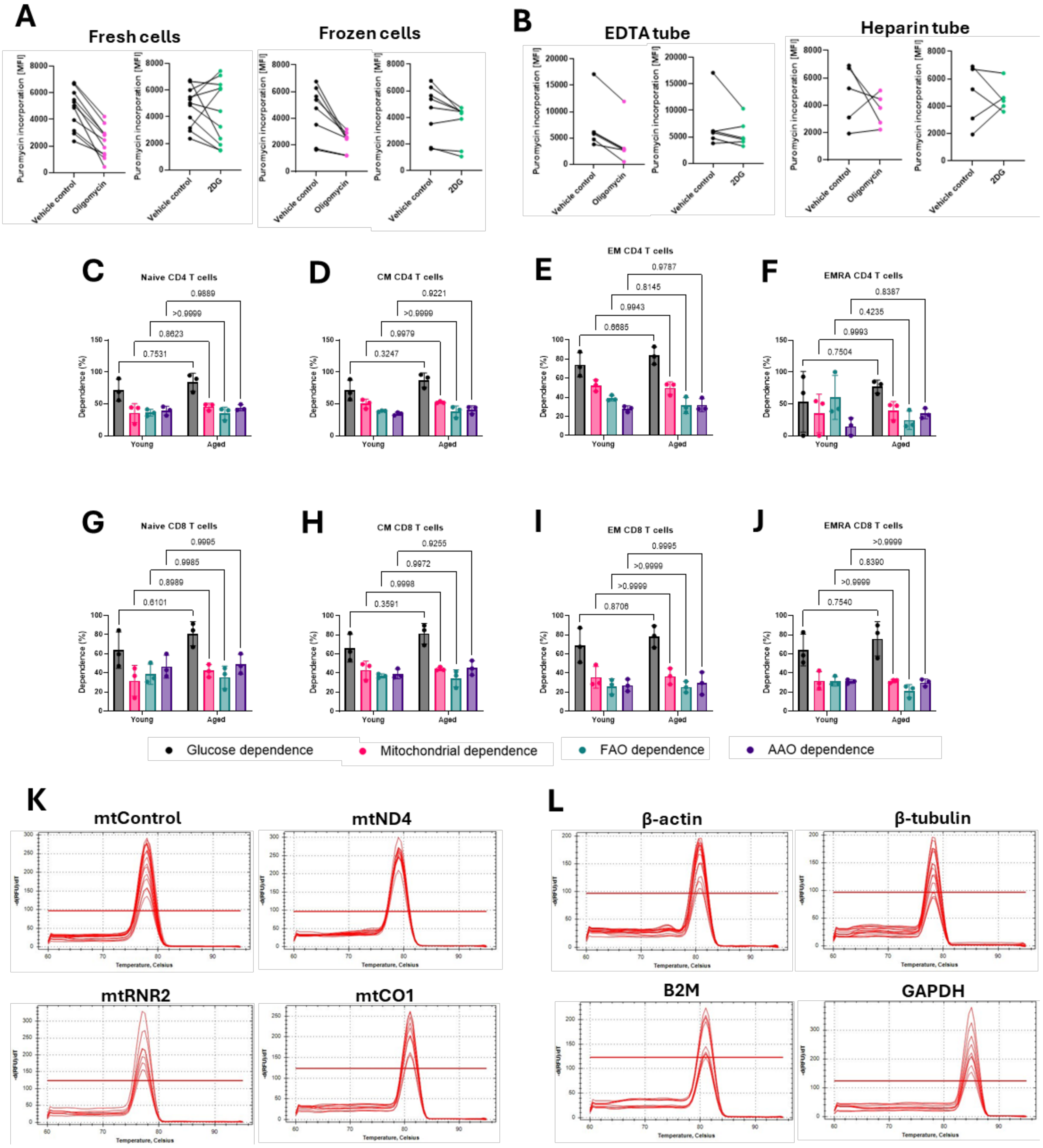

**Supplementary Figure 7. Optimisation of SCENITH and Mitochondrial DNA copy number assays.** Measurement of puromycin incorporation by flow cytometry, at baseline and following inhibition of mitochondrial respiration (oligomycin) and glycolysis (2DG), in **(A)** freshly isolated and frozen peripheral blood mononuclear cells (PBMCs) obtained from young participants (n=8) and **B** matched PBMCs from young participants (n=5) isolated from EDTA or heparin blood tubes. Metabolic dependencies of (**C, G**) Naive, **(D, H)** central memory (CM), **(E, I)** effector memory (EM) and **(F, J)** EMRA (C-F) CD4+ and (G-J) CD8+ T-cell subsets from young (n=3) and aged participants (n=3), established using SCENITH by inhibition of oxidative phosphorylation (oligomycin), glycolysis (2DG), Fatty acid oxidation (Etomoxir) and amino acid oxidation (AAO, BPTES). Representative qRT-PCR melt curve analysis of **K.** mitochondrial and **L.** housekeeping genes from a single young participant, showing a single peak for each primer set, confirming binding specificity. Data are mean ± S.E.M. Statistical significance was determined by Sidak post-test.

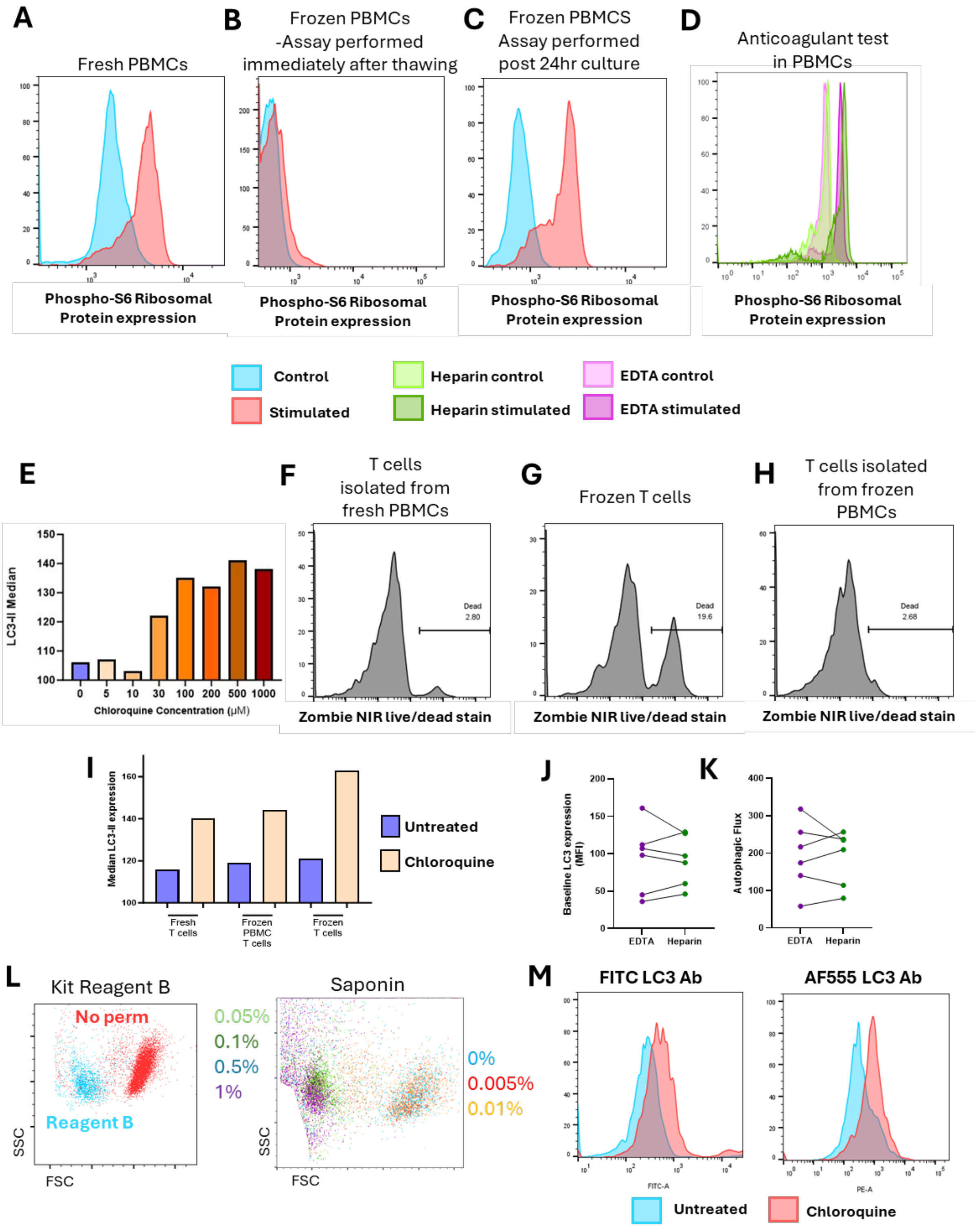


**Supplementary Figure 8. Assessment of T-cell viability, signalling, and autophagy under different processing conditions.** Histograms showing phosphorylated S6 (p-S6) expression at rest and after 30 minutes stimulation with (phorbol 12-myristate 13-acetate) PMA and ionomycin: **(A)** In freshly isolated PBMCs (peripheral blood mononuclear cells), **(B)** PBMCs immediately post-thaw from cryopreservation at –80 °C, **(C)** PBMCs after 24 hours culture post thaw from cryopreservation at –80 °C and **(D)** from PBMCs isolated from heparin- or EDTA blood. **E.** Dose–response of LC3-II (microtubule-associated protein 1 light chain 3-II) expression (mean fluorescence intensity, MFI) following 2 hours incubation with increasing concentrations of chloroquine (CQ) in T cells. Viability of T cells, assessed by Zombie Red staining, **(F)** following fresh isolation from PBMCs, (**G**) from cryopreserved T cells post-thaw, **(H)** and from T cells isolated from frozen PBMCs. **I.** LC3-II expression

(MFI) following 2 hours CQ (100 µM) treatment in freshly isolated T cells, T cells from cryopreserved PBMCs, and directly cryopreserved T cells. **J.** Baseline LC3-II expression and **(K)** autophagic flux in T cells isolated from cryopreserved PBMCs derived from EDTA- or heparin-anticoagulated blood (n = 6). **L.** Forward scatter (FSC) and side scatter (SSC) profiles of T cells following permeabilisation with a commercial kit (left) or increasing concentrations of saponin. **M.** LC3-II expression in untreated and CQ-treated T cells (2 hours, 100 µM), detected with the antibody provided in a commercial autophagy kit versus an Alexa Fluor 555 (AF555)-conjugated anti–LC3-II antibody.

**Supplementary Table 1.** Antibodies used for detection of immune cell populations

| Marker | Fluorophore | Tandem (Y/N) | Supplier |
|---|---|---|---|
| CD3 | PE-Fire 640 | Y | BioLegend (cat. no.: 344860, clone: SK7) |
| CD4 | Spark Blue 550 | N | BioLegend (cat. no.: 344656, clone: SK3) |
| CD45RA | AF700 | N | BioLegend (cat. no.: 304120, clone: HI100) |
| CD25 | APC | N | BioLegend (cat. no.: 302610, clone: BC96) |
| CD57 | mFluor Violet 540 | N | AAT Bioquest (cat. no.: 10570120, clone:HI57a |
| CD28 | Pacific Blue | N | BioLegend (cat. no.: 302928, clone: CD28.2) |
| CD127 | Spark YG 581 | N | BioLegend (cat. no.: 351368, clone: A019D5) |
| LIVE/DEAD | Zombie NIR | N | BioLegend (cat. no.: 423106) |
| CD38 | APC-Fire 810 | Y | BioLegend (cat. no.: 356644, clone: HB-7) |
| CD19 | BB515 | N | BD Biosciences (cat. no.: 564456, clone: HIB19) |
| IgD | BV480 | N | BD Biosciences (cat. no.: 566138, clone: IA6-2) |
| CD27 | BV711 | Y | BioLegend (cat. no.: 302834, clone: O322) |
| CD20 | iFluor532 | N | AAT Bioquest (cat. no.: 10202070, clone: 2H7) |
| CD24 | PE-Alexa 610 | Y | Invitrogen (cat. no.: MHCD2422, clone: SN3) |
| CD8 | PE | N | BioLegend (cat. no.: 303804, clone: QA18A37) |
| CD146 | PerCP-Cy5.5 | Y | BioLegend (cat. no.: 361010, clone: P1H12) |
| CD14 | AF594 | N | BioLegend (cat. no.: 325630, clone: HCD14) |
| HLA-DR | BV570 | Y | BioLegend (cat. no.: 307638, clone: I243) |
| PDGFRa (CD140a) | PE-Cy7 | Y | BioLegend (cat. no.: 323508, clone: 16A1) |
| CD95 | APC-Fire 750 | Y | BioLegend (cat. no.: 305638, clone: DX2) |
| CD31 | BV605 | Y | BioLegend (cat. no.: 303122, clone: WM59) |
| CD69 | PE-Cy5 | Y | BioLegend (cat. no.: 310908, clone: FN50) |
| CD56 | Spark YG 593 | N | BioLegend (cat. no.: 362570, clone: 5.1H11) |
| CD11c | BV421 | N | BioLegend (cat. no.: 301628, clone: 3.9) |
| CD16 | BV510 | N | BioLegend (cat. no.: 302048, clone: 3G8) |
| PD-1 | BV650 | Y | BioLegend (cat. no.: 329950, clone: EH12.2H7) |
| CD90 | BV785 | Y | BioLegend (cat. no.: 328142, clone: 5E10) |
| PDPN | PE-Dazzle 594 | Y | BioLegend (cat. no.: 337028, clone: NC-08) |
| KLRG1 | PE-Fire 810 | Y | BioLegend (cat. no.: 367733, clone: SA231A2) |
| CCR7 | FITC | N | BioLegend (cat. no.: 353216, clone: G043H7) |
| **Intracellular** | | | |
| P21 | CoraLite 594 | N | Proteintech (cat. no.: CL647-10883, clone: Ag1328) |
| P16 | CoraLite Plus 647 | N | Proteintech (cat. no.: CL594-10355, clone: Ag368) |

**Supplementary Table 2.** Markers of immune cell populations for flow cytometry.

| Cell type | Markers |
|---|---|
| **B cells** | $CD16^{-/dim}$ $CD14^{-}$ $CD3^{-}$ $CD19^{+}$ |
| **CD4 T cells** | $CD16^{-/dim}$ $CD14^{-}$ $CD3^{+}$ $CD4^{+}$ |
| **CD8 T cells** | $CD16^{-/dim}$ $CD14^{-}$ $CD3^{+}$ $CD8^{+}$ |
| **Central memory CD4 T cells** | $CD16^{-/dim}$ $CD14^{-}$ $CD3^{+}$ $CD4^{+}$ $CCR7^{+}$ $CD45RA^{-}$ |
| **Central memory CD8 T cells** | $CD16^{-/dim}$ $CD14^{-}$ $CD3^{+}$ $CD8^{+}$ $CCR7^{+}$ $CD45RA^{-}$ |
| **Circulation endothelial cells** | $CD16^{-/dim}$ $CD14^{-}$ $CD3^{-}$ $CD19^{-}$ $CD45RA^{-}$ $PDPN^{-}$ $CD146^{-}$ $CD31^{+}$ |
| **Classical monocytes** | $CD16^{-/dim}$ $CD3^{-}$ $CD19^{-}$ $CD14^{+}$ $HLA\text{-}DR^{+/-}$ $CD16^{-}$ |
| **Dendritic cells** | $CD16^{-/dim}$ $CD14^{-}$ $CD3^{-}$ $CD19^{-}$ $CD11c^{+}$ $HLA\text{-}DR^{+}$ |
| **Double negative B cells** | $CD16^{-/dim}$ $CD14^{-}$ $CD3^{-}$ $CD19^{+}$ $IgD^{-}$ $CD27^{-}$ |
| **Early natural killer cells** | $CD16^{-/dim}$ $CD14^{-}$ $CD3^{-}$ $CD19^{-}$ $CD11c^{-}$ $HLA\text{-}DR^{-}$ $CD16^{-}$ $CD56^{+}$ |
| **Effector memory CD4 T cells** | $CD16^{-/dim}$ $CD14^{-}$ $CD3^{+}$ $CD4^{+}$ $CCR7^{-}$ $CD45RA^{-}$ |
| **Effector memory CD8 T cells** | $CD16^{-/dim}$ $CD14^{-}$ $CD3^{+}$ $CD8^{+}$ $CCR7^{-}$ $CD45RA^{-}$ |
| **EMRA CD4 T cells** | $CD16^{-/dim}$ $CD14^{-}$ $CD3^{+}$ $CD4^{+}$ $CCR7^{-}$ $CD45RA^{+}$ |
| **EMRA CD8 T cells** | $CD16^{-/dim}$ $CD14^{-}$ $CD3^{+}$ $CD8^{+}$ $CCR7^{-}$ $CD45RA^{+}$ |
| **Intermediate monocytes** | $CD16^{-/dim}$ $CD3^{-}$ $CD19^{-}$ $CD14^{+}$ $HLA\text{-}DR^{+/-}$ $CD16^{+}$ |
| **Mature natural killer cells** | $CD16^{-/dim}$ $CD14^{-}$ $CD3^{-}$ $CD19^{-}$ $CD11c^{-}$ $HLA\text{-}DR^{-}$ $CD16^{+}$ $CD56^{+}$ |
| **Memory B cells** | $CD16^{-/dim}$ $CD14^{-}$ $CD3^{-}$ $CD19^{+}$ CD38- $CD24^{+}$ |
| **Monocyte** | $CD16^{-/dim}$ $CD3^{-}$ $CD19^{-}$ $CD14^{+/-}$ $HLA\text{-}DR^{+/-}$ |
| **Naïve B cells** | $CD16^{-/dim}$ $CD14^{-}$ $CD3^{-}$ $CD19^{+}$ $IgD^{+}$ $CD27^{-}$ |
| **Naïve CD4 T cells** | $CD16^{-/dim}$ $CD14^{-}$ $CD3^{+}$ $CD4^{+}$ $CCR7^{+}$ $CD45RA^{+}$ |
| **Naïve CD8 T cells** | $CD16^{-/dim}$ $CD14^{-}$ $CD3^{+}$ $CD8^{+}$ $CCR7^{+}$ $CD45RA^{+}$ |
| **Natural killer cells** | $CD16^{-/dim}$ $CD14^{-}$ $CD3^{-}$ $CD19^{-}$ $CD11c^{-}$ $HLA\text{-}DR^{-}$ |
| **Non-classical monocytes** | $CD16^{-/dim}$ $CD3^{-}$ $CD19^{-}$ $CD14^{-}$ $HLA\text{-}DR^{+/-}$ $CD16^{+}$ |
| **Non-switched memory B cells** | $CD16^{-/dim}$ $CD14^{-}$ $CD3^{-}$ $CD19^{+}$ $IgD^{+}$ $CD27^{+}$ |
| **Plasmablasts** | $CD16^{-/dim}$ $CD14^{-}$ $CD3^{-}$ $CD19^{+}$ $CD38^{+}$ $CD24^{-}$ |
| **Regulatory B cells** | $CD16^{-/dim}$ $CD14^{-}$ $CD3^{-}$ $CD19^{+}$ $CD38^{+}$ $CD24^{+}$ |
| **Regulatory T cells** | $CD16^{-/dim}$ $CD14^{-}$ $CD3^{+}$ $CD4^{+}$ $CD25^{+}$ |
| **Switched memory B cells** | $CD16^{-/dim}$ $CD14^{-}$ $CD3^{-}$ $CD19^{+}$ $IgD^{-}$ $CD27^{+}$ |
| **T cells** | $CD16^{-/dim}$ $CD14^{-}$ $CD3^{+}$ |
| **Terminal natural killer cells** | $CD16^{-/dim}$ $CD14^{-}$ $CD3^{-}$ $CD19^{-}$ $CD11c^{-}$ $HLA\text{-}DR^{-}$ $CD16^{+}$ $CD56^{-}$ |

**Supplementary Table 3.** Primer sequences used for measurement of mtDNA copy number.

| **mtRNR2** | **β-actin** |
|---|---|
| Forward primer: CACCCAAGAACAGGGTTTGT<br>Reverse primer: TGGCCATGGGTATGTTGTTA | Forward primer: AAGTCCTGCCCTCATTTCCC<br>Reverse primer: GGATGTCCACGTCACACTTCA |
| **mtND4** | **β-tubulin** |
| Forward primer: CTGTTCCCCAACCTTTTCCT<br>Reverse primer: CCATGATTGTGAGGGGTAGG | Forward primer: TTTGGGTGCAAGGACACAGC<br>Reverse primer: CTCCCCCTACTGCCCCATAA |
| **mtCO1** | **B2M** |
| Forward primer: ATACCAAACGCCCCTCTTCG<br>Reverse primer: TGTTGAGGTTGCGGTCTGTT | Forward primer: TGCTGTCTCCATGTTTGATGTATCT<br>Reverse primer: TCTCTGCTCCCCACCTCTAAGT |
| **mtControl region** | **GAPDH** |
| Forward primer: CTAAATAGCCCACACGTTCCC<br>Reverse primer: AGAGCTCCCGTGAGTGGTTA | Forward primer: CTATGGACACGCTCCCCTGA<br>Reverse primer: AGGAGCCAGTCTTGGATGAGA |

**Supplementary Table 4.** Constituents of RCP reactions performed to measure the mtDNA copy number.

| Reagent | Amount per reaction | Specifications |
|---|---|---|
| PowerUp™ SYBR™ Green Master Mix | 5 µl | A25742, Thermo Fisher, UK |
| Forward primer (10 µM solution) | 1 µl | 41105327, Merck, UK |
| Reverse primer (10 µM solution) | 1 µl | 41105327, Merck, UK |
| DNA template (6 ng/µl dilution) | 2 µl | N/A |
| Nuclease-free water | 1 µ | AM9920, Thermo Fisher, UK |

**Supplementary Table 5.** Settings used to perform the RCP reactions measuring the mtDNA copy number.

| Step | Temperature | Duration | Number of cycles |
|---|---|---|---|
| DNA polymerase activation | 50 °C | 2 min | N/A |
| Initial denaturation | 95 °C | 2 min | N/A |
| Denaturation<br>Annealing and extension | 95 °C<br>61 °C | 15 sec<br>30 sec | 40 cycles |
| Melt curve | 60-95 °C | | N/A |
